# Tetraspanin CD81 promotes leukemia stem cell function and represents a new therapeutic vulnerability in acute myeloid leukemia

**DOI:** 10.1101/2023.09.20.558656

**Authors:** Fanny Gonzales, Pauline Peyrouze, Thomas Boyer, Soizic Guihard, Francois Sevrin, Djohana Laurent, Adriana Plesa, Adeline Barthelemy, Antonino Bongiovanni, Nicolas Pottier, Claude Preudhomme, Nicolas Duployez, Céline Berthon, Christophe Roumier, Meyling Cheok

## Abstract

Despite important progress over the last decade, acute myeloid leukemia (AML) is still associated with poor clinical outcome. Novel potent therapies ideally effective against AML stem cells (LSC), a major driver of leukemia initiation and progression, are urgently needed. In particular, targeting common AML-associated antigens at the stem and progenitor cell level represents an attractive therapeutic strategy to achieve deep long-term remissions and is currently the subject of intensive research efforts. In this study, we identified the tetraspanin CD81, a cell surface antigen frequently expressed on AML cells including LSC, as a new determinant of relapse and poor prognosis. CD81 expression was higher in AML cells compared to normal bone marrow cells, and more markedly expressed at relapse. We further showed that modulation of CD81 expression using gain- and loss-of-function approaches affected leukemia aggressiveness, tumor burden, LSC-homing and - xenoengraftment as well as mouse survival. Finally, anti-hCD81 monoclonal antibody-treatment combined with standard chemotherapy in mice with pre-established AML not only reduced leukemia burden but also prolonged relapse-free and overall survival. Collectively, these results identified a new efficacious and safe pharmacological strategy for targeting LSC, opening up novel therapeutic avenues to improve AML outcome.

**Key points:**

- CD81 expression in AML including LSC is a new determinant of aggressive disease and poor prognosis.
- Anti-hCD81 monoclonal antibody-treatment of AML xenografts reduced leukemia burden and improved survival rates.

## Introduction

Acute myeloid leukemia (AML) is the most common acute leukemia in adults with an annual incidence of approximately 3 to 4 cases per 100,000 persons worldwide.^1^ Over the past few decades, the therapeutic strategy has mostly relied on intensive combination chemotherapy, notably cytarabine and anthracyclines, often followed by hematopoietic stem cell transplantation (HSCT). More recently, various targeted therapies have become available for specific patients,^2^ but despite this progress, mortality in AML patients remains high and poor clinical outcome is mainly due to disease relapse or treatment failure, occurring in up to 40% of younger and 80% of older adult AML cases.^3–6^ Even though several relapse risk factors are known,^7,8^ a better understanding of the mechanisms leading to refractory or relapsed AML is still needed to develop more potent therapeutic strategies.

Therapy-resistant and quiescent leukemia stem cells (LSC), present within the leukemia bulk at very low frequency and functionally defined as capable to initiate AML when transplanted into immunocompromised mice,^9^ are considered to be the major cause of disease progression and relapse.^10^ Development of new potent treatment strategies to eradicate LSC has thus recently become the focus of intensive research.^11^ While various distinct cell surface proteins such as CD33,^12^ CD123,^13^ CD44, CD47,^14,15^ TIM-3,^16^ CLL-1^17,18^ and CD45RA^19^ have been found enriched or absent such as CD90^20^ on LSC, therapies targeting these markers are lagging behind expectations, mainly due to their overlapping expression with hematopoietic stem and progenitor cells (HSPC) as well as the lack of antigens broadly and consistently expressed on LSC in most AML subtypes. Therefore, the identification of new druggable and discriminating LSC markers remains essential for the successful development of a selective anti-LSC therapy.

CD81 refers to a cell surface protein, also known as “target of an antiproliferative antibody 1” (TAPA-1),^21^ that belongs to the evolutionary conserved tetraspanin superfamily of proteins. CD81 is the structural component of specialized membrane microdomains known as tetraspanin-enriched microdomains,^22–24^ and plays an important role in signal transduction by interacting with various proteins such as integrins and several distinct membrane proteins.^25,26^ In particular, CD81 has been shown to regulate B cell signaling by interacting with CD19 and CD21 on B-lymphocytes^27,28^ and is known to be overexpressed in several hematological malignancies implicating B cells such as lymphoma^21,29^ and acute lymphoblastic leukemia.^30^ Importantly, previous studies have demonstrated that immunotargeting of CD81 represents a new effective treatment for B cell lymphoma.^29,31^ Nevertheless, despite its adverse role in B cell malignancies, its functional implication in AML remains to be characterized.

In this study, using a combination of *in vitro* and *in vivo* approaches, we demonstrated that CD81 is an important determinant of AML progression, drug resistance and leukemia stemness. Finally, we provide proof of concept that targeting CD81 may represent a safe and efficacious therapeutic strategy to eradicate LSC in AML.

## Materials and Methods

### Patients and cell line models

In this study, we included 11 normal bone marrow (BM) samples and 290 primary AML including 38 relapse samples recruited at Lille University Hospital (2009-2016, Table 1). All patients have signed the informed consent form and the Institutional Review Board approved these studies (ANA-FE-BMH 20221007). CD81 protein expression (*N* = 290, Supplemental Figure 1), CD81 mRNA expression (*N* = 53), LSC17 gene expression score (*N* = 107) and AML xenoengraftment (*N* = 59) were performed as described previously.^32–34^ Diagnostic patient characteristics are given in Table 1, which excluded not intensively treated (*N* =48), pediatric (*N* = 16) and patients with incomplete clinical data (*N* = 8).

**Table 1.** Diagnostic parameters of the study cohort treated with standard chemotherapy.

| Parameter | Total | CD81 low | CD81 high | P-value |
| --- | --- | --- | --- | --- |
| No. of patients (%) | 150 (100) | 68 (45) | 82 (55) | - |
| Age [y], median (min-max) | 55.6 (17.3-77.6) | 53.6 (17.3-77.6) | 56.3 (17.8-75.7) | 0.71 |
| Gender (M/F) | 72/78 | 31/37 | 41/41 | 1.0 |
| WBC [$\times 10^9/L$], median (min-max) | 27.2 (1-405) | 13.4 (1-249) | 46.1 (1-405) | 0.010* |
| BM blast [%], median (min-max) | 70 (5-98) | 61.5 (5-97) | 72 (5-98) | 0.114 |
| **Cytogenetics** |  |  |  | 0.86 |
| Favorable, N (%) | - | - | - |  |
| Normal, N (%) | 85 (56.7) | 41 (60.3) | 44 (53.6) |  |
| Abnormal, N (%) | 35 (23.3) | 13 (19.1) | 22 (26.8) |  |
| Adverse, N (%) | 26 (17.3) | 12 (17.6) | 14 (17.1) |  |
| NA, N (%) | 4 (2.7) | 2 (3) | 2 (2.5) |  |
| **ELN2022 risk** |  |  |  | 0.73 |
| Favorable, N (%) | 26 (17.3) | 15 (22.0) | 11 (13.0) |  |
| Intermediate, N (%) | 94 (62.7) | 39 (57.4) | 55 (67.1) |  |
| Adverse, N (%) | 26 (17.3) | 12 (17.6) | 14 (17.1) |  |
| NA, N (%) | 4 (2.7) | 2 (3) | 2 (2.5) |  |

\* $P < 0.05$

AML cell lines U-937, OCI-AML3, HNT-34 (DSMZ), and BM stromal fibroblasts HS-5 (ATCC) were cultured as recommended. Authenticated AML cell lines (FTA Kit, LGC, ATCC-135-XV-20) were transduced using the luciferase containing lentiviral vector (pLenti PGK V5-LUC Neo (623-2), Addgene). U-937 cells were transfected with pCDM8 hCD81^21^ (Addgene) using the Amaxa Nucleofactor II, (kit C, W001; Lonza) and CD81 positive cells sorted (ARIA III BD, BD Biosciences). HNT-34 and OCI-AML3 cells were transduced with shRNA-CD81 or non-targeting (NT) shRNA lentiviral vectors (TRCN0000300291[sh291], TRCN0000300293[sh293], TRCN0000300433[sh433] or TRC2 pLKO.5-puro; Sigma-Aldrich).

### AML xenografts

Patient-derived xenografts (PDX) and cell-line-derived xenografts (CDX) were generated using NSG immunodeficient mice as previously described^33^ (Supplemental Figure 2). We monitored CDX engraftment by *in vivo* imaging (luciferin 0.1 mg/kg IP, Xenogen IVIS 50, Perkin Elmer). We collected femurs and sternums from CDX when hCD45 was >1% in peripheral blood (PB) and from control mice injected with PBS. Bones were fixed with 4% paraformaldehyde-PBS overnight at 4°C, decalcified (DC1, Q Path) for 18h at 4°C, dehydrated and embedded in paraffin. Immunohistochemistry was performed on 4 μm sections using the BenchmarkUltra (Roche), anti-hCD45 (M0701, Dako) and UltraView DAB Detection Kit (Ventana). We evaluated AML cell homing by flow cytometry (FCM) analysis of BM using a minimum of 1 x10^6^ acquired events, 72h after IV injection of 12 x10^6^ (OCI-AML3) or 20 x10^6^ (U-937) AML cells. French ethics committee approved all animal experiments (APAFIS#33794-2021100815558055).

### CD81 expression

CD81 cell surface protein expression was determined by FCM (LSR Fortessa X20, BD Biosciences), 5 x10^5^ cells were labeled with 1 μg anti-hCD81-PE clone 5A6 or 1 µg isotype control-PE (349506, 400114; Biolegend)^32^ and by Western blotting (4-12% SDS-PAGE, clone JS-81, BD Biosciences). The FCM antibody panel and gating strategy are detailed in Supplemental Figure 1.^33^ CD81 mRNA was quantified by Taqman RT-PCR and GAPDH normalized (CD81: Hs00174717_m1, GAPDH: Hs03929097_g1, StepOnePlus Real-Time PCR System, ThermoFisher).

### *In vitro* drug resistance

We assessed drug resistance *in vitro* as previously described.^35^ Drug concentration ranges were 0.000042-3.33 µg/mL (daunorubicin [DNR]) and 0.0025-40 µg/mL (cytarabine [AraC]) and drug exposure times were 72h (AML cell lines) and 96h (primary AML cells).

### Cell adhesion, migration and invasion

We coated 96-well plates with fibronectin (1 μg per cm^2^, S5171, Sigma-Aldrich) or seeded 4 x10^4^ HS-5 for 48h (70-80% confluence). Then, cells in 100 μL (U-937, OCI-AML3 [1 x10^6^]; HNT-34 [0.5 x10^6^]) were allowed to adhere in triplicates for 90 min (U-937, OCI-AML3) or 30 min (HNT-34). Plates were washed three times with PBS and bioluminescence measured (luciferin 0.3 mg/mL, Xenogen IVIS 50; Perkin Elmer). We used 24-well transwell inserts and plates (5 μm 29442-118, Corning) to evaluate cell migration and invasion. Lower chambers were seeded with 5 x10^5^ HS-5 in 300 μL 24h prior to the experiment. Then, 3 x10^5^ AML cells in 300 μL were loaded onto the upper chambers and cultured for 24h. Cells migrated to the lower chambers were quantified by bioluminescence as described above. For invasion experiments, the upper chambers were loaded with a gel matrix using 100 µL Basement Membrane Extract (BME) at 0.1X per insert (Cultrex 5X BME Solution, Trevigen) 4h prior to AML cell loading. All experiments were performed in duplicates. Adhesion, migration and invasion rates were determined as the percentage of cells *vs.* input control (Supplemental Figure 3).

### Confocal microscopy

AML cells (6 x10^4^ cells/cm^2^) were seeded for 24h on poly-D-Lysine (Gibco) coated Lab-Tek II Chamber Slides (Nunc) at a density of. Then, slides were rinsed with PBS and cells fixed with 4% paraformaldehyde-PBS for 20 min and permeabilized with PBS-Triton X-100 0.1% for 10 min at room temperature (RT). After blocking with PBS-BSA 2% for 45 min, cells were stained with Alexa Fluor 488 phalloidin (1:40, Invitrogen) for 30 min at RT in the dark, and nuclei were stained with DAPI (ThermoFisher) for 10 min. We generated confocal microscopy images (LSM710, Zeiss) and estimated cellular diameter and circularity using ImageJ, (https://imagej.nih.gov/ij/)^36^.

### Anti-hCD81 antibody-treatment

Cells were exposed to anti-hCD81 monoclonal antibody or isotype control for 4h with 1 µg per 1 x10^6^ cells in 100 µL. AML bearing mice were treated with anti-hCD81 antibody *vs.* isotype control (JS-81, BD Biosciences) 100 µg, IP using three or five doses in OCI-AML3 CDX and PDX, respectively in combination with chemotherapy.^37^ *In vivo* imaging of OCI-AML3 CDX and FCM of PDX blood^33^, was used to track leukemia engraftment and disease progression.

### Colony-Forming Unit (CFU) assays

CFU assays were performed according to the MethoCult™ H4034 Optimum protocol (Stemcell Technologies). Briefly, 5 x10^4^ and 2.5 x10^4^ human BM mononuclear cells (BMNC, Lonza) were plated in triplicate and cultured for 10 days. Colonies were counted and normalized to the number of cells seeded on day 0.

### Cell cycle and viability analyses

Cell cycling of BMNC was determined on 0.25 x10^6^ BMNC washed in PBS and fixed in cold ethanol 70% at -20°C for 2h. Then, cells were washed twice in PBS-FBS 1% and stained with 20 µL of anti Ki-67 (FITC Mouse Anti-Ki-67 Set, BD Biosciences) for 30 min at RT in the dark. Cells were washed in PBS-FBS 1%, incubated with RNaseA 2 µg/mL for 15 min at 4°C and stained with PI 35 µg/mL for 30 min before analysis by FCM. Viability of BMNC (0.25 x10^6^) was assessed by Annexin V and 7-AAD staining (APC Annexin V Apoptosis Detection Kit 7-AAD, Biolegend) using FCM.

### Statistics

We used Student’s t-test and Pearson correlation statistics unless otherwise specified and results included sample means, standard deviation and *R*^2^. We evaluated time-to-xenoengraftment using Cox proportional hazard model with *P*-values based on the log-rank test. We censored time at transplantation date in the event of HSCT or at the date of last follow-up and defined overall survival (OS) as the time from date of inclusion to date of death from any cause. Relapse-free survival (RFS) was determined in patients achieving complete remission (CR) and is defined as the time from date of CR to date of first relapse, or in the case of no relapse, time was censored at the date of last follow-up. The alpha level was set to 5% and all statistical tests were two-sided. We used GraphPad Software version 10 (Prism). Graphs and analyses were computed using Kaplan-Meier or Cumulative incidence plots, hazard ratio (HR) and 95% confidence intervals (CI) were determined using R-v.4.2.0.^38^ CD81 mRNA expression and survival of publicly available Beat-AML dataset^39^ were obtained *via* the cBioPortal.^40,41^ Similar to our cohort the analyses included AML (non-FAB-M3, non-core-binding factor (CBF), age >50, chemotherapy).

## Results

### CD81 overexpression is associated with AML relapse and poor prognosis

Previously, we determined that high CD81 expression prognostic using a cohort of 134 diagnostic AML patients.^32^ These initial findings are now extended using a cohort of 252 diagnostic and 38 relapse samples. Since CBF-AML have well-known favorable prognosis, we first compared CD81 protein expression between CBF and non-CBF AML and found it to be lower in the CBF group (*N* = 28, Figure 1A). Therefore, we excluded CBF-AML to avoid selection bias and confirmed that higher CD81 protein expression adversely affected RFS and OS (*N* = 150, RFS: HR [CI] = 1.91 [1.12-3.28]; OS: HR [CI] = 1.88 [1.07-3.32]; Figure 1B). Interestingly, we also showed that higher CD81 protein expression negatively affected OS within *NPM1* mutated AML, a molecular subtype with heterogeneous clinical response (*N* = 54, OS: HR [CI] = 7.71 [1.00-59.23]; Figure 1C), independently of co-existing *FLT3-ITD* mutational status (data not shown) and a similar trend was obtained for relapse (RFS: *P* = 0.15; Figure 1C). As both CD81 mRNA expression and protein abundance were correlated in our cohort (*P* = 0.01, Figure 1D), we further characterized the detrimental effect of CD81 overexpression on AML prognosis independently using the publicly available mRNA data from the Beat AML dataset.^39^ Consistent with our results, CBF-AML from the Beat-AML cohort also showed a significant lower CD81 mRNA expression compared to non-CBF AML (Figure 1E). Importantly, we confirmed the negative impact of high CD81 mRNA on OS not only within non-CBF AML but also within *NPM1* mutated AML (non-CBF AML: HR [CI] = 1.80 [1.00-3.26]; *NPM1* mutated AML: HR [CI] = 5.39 [1.58-18.36]; Figure 1F). Finally, given the prognostic effect of high CD81 expression, we hypothesized that blasts at diagnosis exhibit an increased expression of this protein and an even higher CD81 expression at relapse, a disease state usually refractory to treatment. As expected, CD81 was overexpressed in AML compared to BM obtained from healthy individuals (Figure 1G) and AML at relapse exhibited higher CD81 expression when compared to samples at diagnosis (Figure 1H) or using paired diagnostic-relapsed samples (67% *vs.* 19%, Figure 1I).

**Figure 1.**
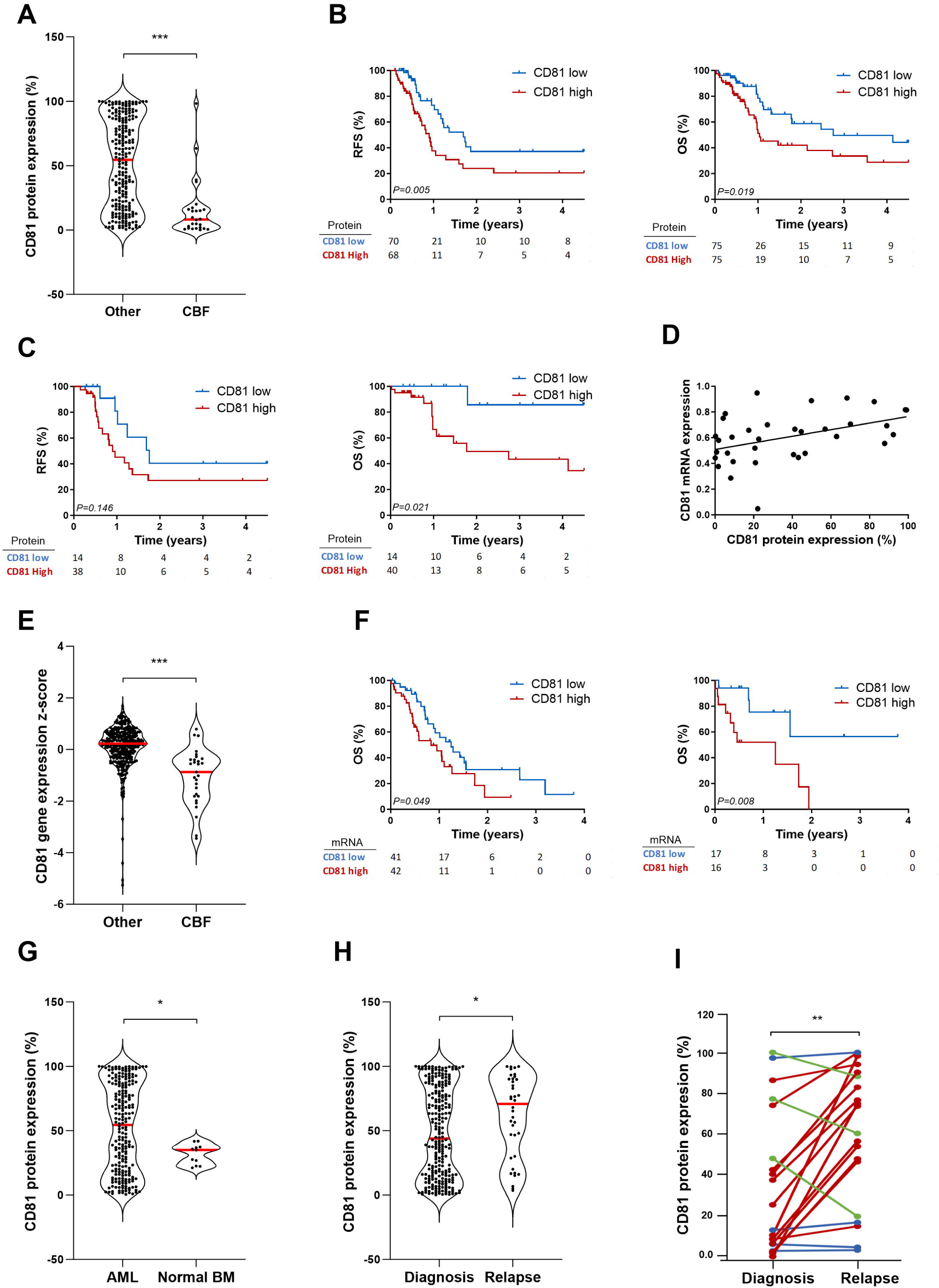
CD81 expression is associated with AML relapse and poor clinical outcome. (A) Violin plot comparing CD81 expression in non-CBF *vs*. CBF-AML (*N* = 252). (B) Kaplan-Meier curves showing worse RFS and OS of patients with higher CD81 expression in their AML and similarly, (C) in a subset of patients with *NPM1* mutant AML. (D) Scatterplot correlating CD81 mRNA and protein expression (*N* = 33, *R*^2^ = 0.19, *P* = 0.019). (E) Violin plot depicting that favorable CBF AML expressed lower levels of CD81 mRNA expression (Beat AML dataset, *N* = 364). (F) Kaplan-Meier curves of patients stratified by median CD81 mRNA at AML diagnosis, showing worse OS for patients with AML overexpressing CD81 overall (left panel) and within *NPM1* mutant AML (right panel). (G) Violin plot indicating CD81 overexpression in AML compared to normal BMNC, (H) which increased at relapse compared to leukemia diagnosis on all AML (*N* = 290) and (I) also in paired AML samples (red: increase, blue: no change, green: decrease; *N* = 19; paired t-test). \**P* <0.05, \*\**P* <0.005, \*\*\**P* <0.0005.

Collectively, these findings demonstrated that CD81 overexpression on AML blasts adversely affects AML outcome and may contribute to the aggressiveness characterizing blasts at relapse.

### *In vitro* modulation of CD81 expression affects xenoengraftment and leukemia aggressiveness

To further strengthen the deleterious role of CD81 in AML, we modulated its expression in human AML cell lines: OCI-AML3, HNT-34 and U-937. Overexpression of CD81 was performed in U-937 cells, which have undetectable CD81 expression, whereas silencing of CD81 was achieved using a shRNA approach in HNT-34 and OCI-AML3 cells, which by contrast highly express CD81 (Supplemental Figure 4). Since we linked primary CD81 overexpression to AML relapse, we first assessed whether modulating CD81 expression affected chemoresistance. We found that U937-CD81 overexpressing cells were more resistant to daunorubicin, whereas CD81-depleted cells (OCI-AML3, HNT-34) were more sensitive to daunorubicin and cytarabine (Figure 2A). Then, we tested engraftment efficacy, a recognized indicator of AML aggressiveness,^42,43^ evaluated by *in vivo* imaging of leukemia cells exhibiting either high or low CD81 expression in immunocompromised NSG mice. As shown in Figure 2B, xenoengraftment was more successful in U-937 overexpressing CD81 compared to control cells, whereas CD81-depleted OCI-AML3 cells exhibited reduced xenoengraftment capacity. Consistent with this, AML cell homing was similarly affected by CD81 modulation (Figure 2C). As AML engraftment implicates multiple cell-based processes such as adhesion, migration and invasion,^44^ we also examined the contribution of CD81 on each of these cellular programs (Figure 3A). Of particular interest, overexpression of CD81 in the U-937 model increased AML cell adhesion using fibronectin-coated plates or HS-5 fibroblast co-cultures, whereas CD81 depletion in both OCI-AML3 and HNT-34 cell models had the opposite effect. Similar results were obtained for cellular migration and invasion of AML cells into a semisolid gel matrix. Confocal microscopy using phalloidin staining additionally showed that CD81 overexpressing AML cells formed filopodia-like membrane protrusions, a typical morphological change reminiscent of the cellular processes investigated (Figure 3B). Finally, we evaluated whether CD81 modulation also affected leukemia aggressiveness by assessing mouse survival and AML infiltration into various tissues using hCD45 as a marker of leukemic cells. As shown in Figure 4A, mice engrafted with U937-CD81 overexpressing cells have higher leukemia burden in blood as well as in BM and its surrounding muscle tissue (Figure 4B) along with a worse survival (HR [CI] = 3.42 [1.32-9.90], Figure 4C). By contrast, CD81-depleted cells induced a less aggressive phenotype with reduced tissue infiltration (OCI-AML3, Figure 4D) and a better survival (HR [CI] = 0.33 [0.12-0.09], Figure 4E).

**Figure 2.**
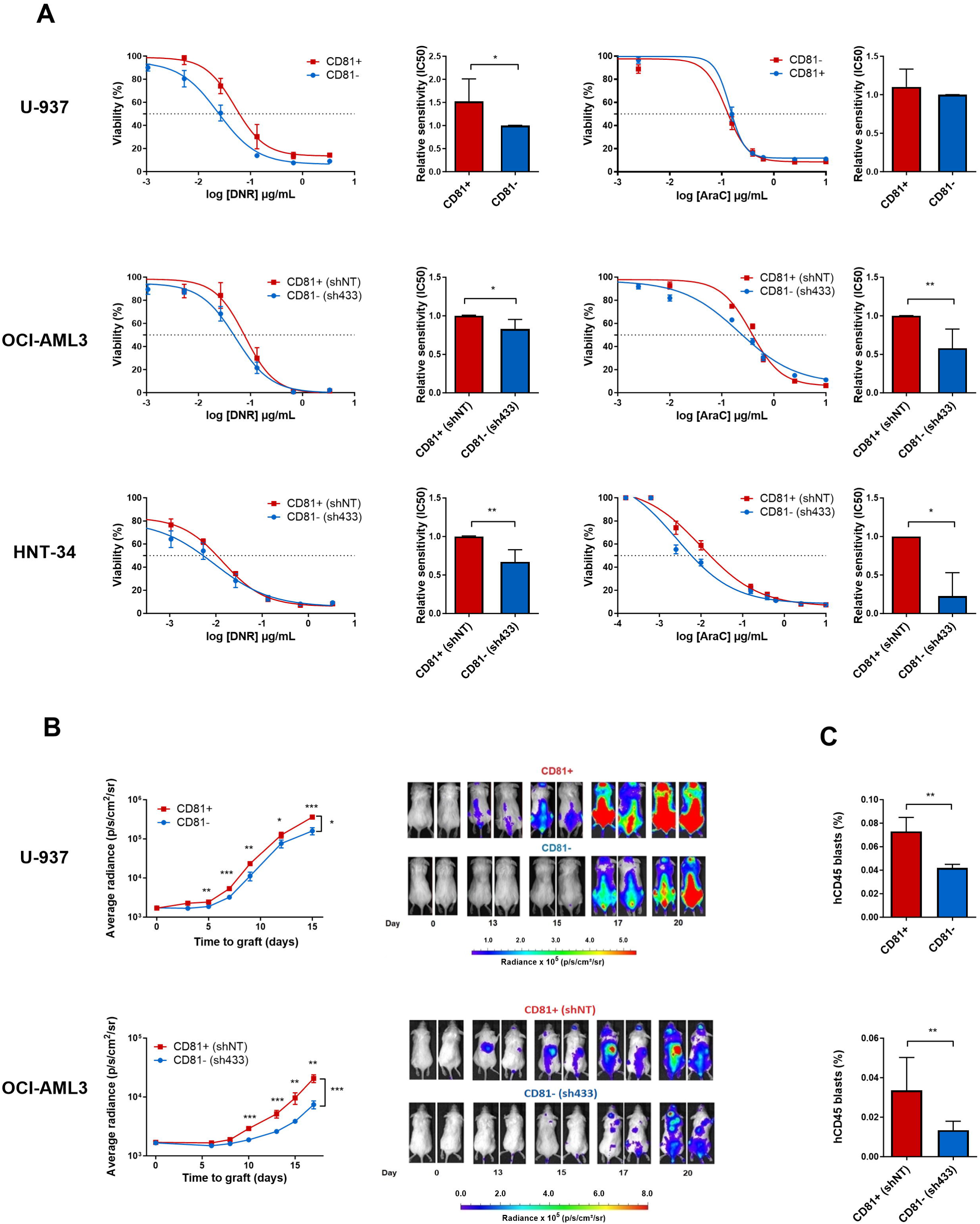
AML models of high CD81 expression are chemoresistant and capable to engraft in immunodeficient mice. (A) Daunorubicin (DNR, left) and cytarabine (AraC, right) drug-response curves and bar charts of IC_50_ using AML models with either CD81 overexpression (red, CD81+) or silenced CD81 (blue, CD81-) *vs.* respective controls U-937 [top], OCI-AML3 [middle], HNT-34 [bottom] evaluated by *in vitro* drug resistance (*N* >3). (B) Line graphs of luminescence and *in vivo* images of xenoengrafts demonstrating increased tumor burden of U-937 CD81 overexpressing (CD81+, red) cells compared to control (*N* =17, Supplemental Figure 8A). Conversely, OCI-AML3 CD81 silencing (CD81-, blue) decreased tumor burden compared to control (*N* =11, Supplemental Figure 8B). (C) AML cell homing by FCM analysis of BM cells 72h after IV injection indicate increased BM homing when CD81 was overexpressed (top panel, U-937, *N =* 3, MWU) whereas it decreased when CD81 was silenced (bottom panel, OCI-AML3, *N =* 5, MWU). \**P* <0.05, \*\**P* <0.005, \*\*\**P* <0.0005.

**Figure 3.**
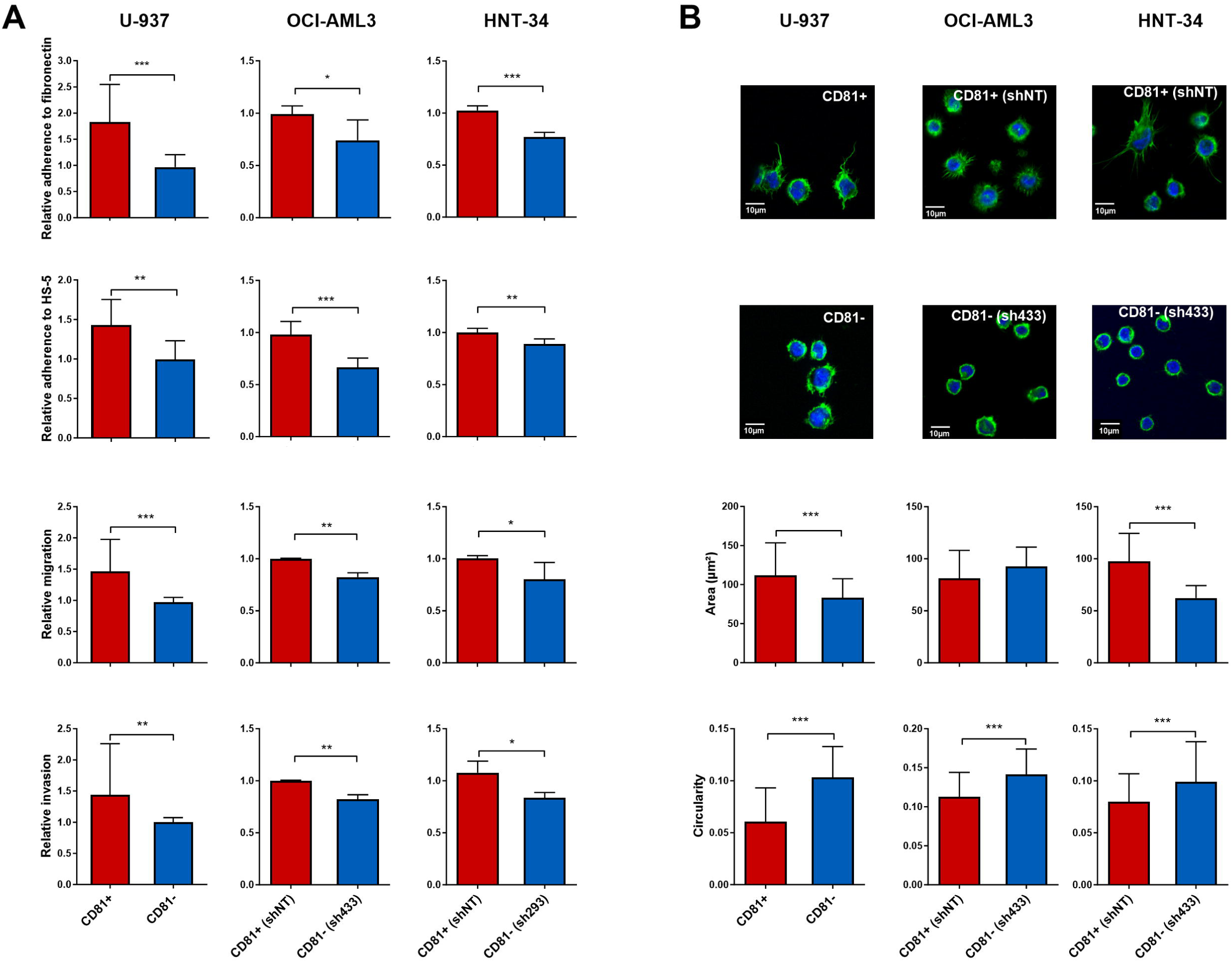
CD81 overexpression modeling in AML reveals enhanced adhesion, migration and invasion capacity and filopodia-like membrane protrusion formation. CD81 overexpression (CD81+, red) induced cell adhesion to fibronectin and HS-5 fibroblasts, migration and invasion compared to control (U-937, *N* >4, left panels). Conversely, CD81 silencing (CD81-, blue) decreased cell adhesion in both OCI-AML3 (*N* >3, middle panel) and HNT-34 (*N* >3, right panel) compared to control. (B) Confocal microscopy displaying filopodia-like cell membrane protrusions in high CD81 expressing AML models (top images, U-937, OCI-AML3, HNT-34) compared to their CD81 low expression counterparts exempt of membrane protrusions (bottom images). Corresponding bar charts summarize cell size and circularity detected by confocal microscopy (U-937, *N* = 3, 218/434 cells; OCI-AML3, *N* = 2, 295/170 cells; HNT-34, *N* = 2, 263/173 cells). \**P* <0.05, \*\**P* <0.005, \*\*\**P* <0.0005.

**Figure 4.**
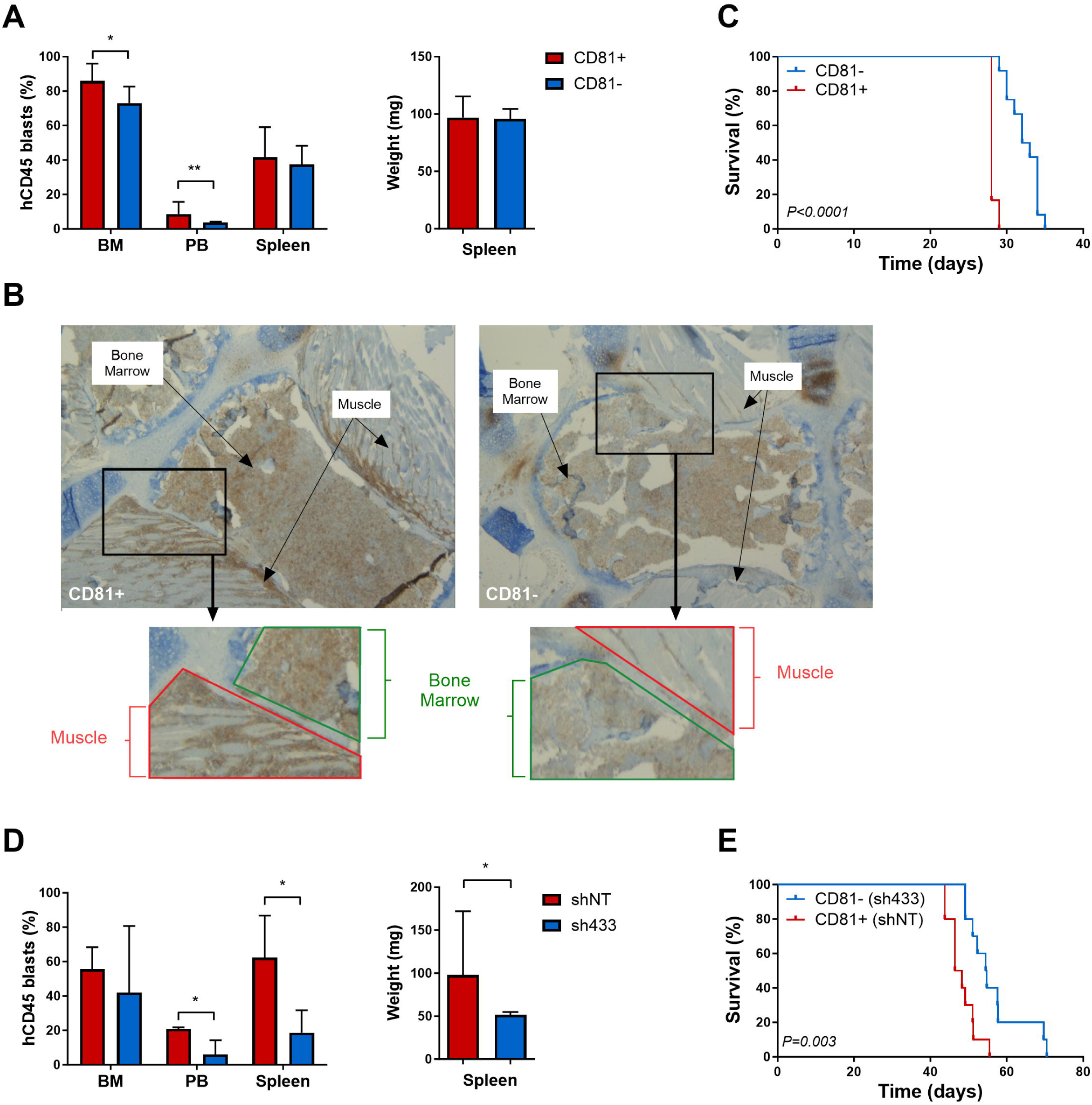
CD81 overexpression models of AML induce leukemia burden and aggressive disease in xenoengrafted mice. (A) CD81 overexpression (CD81+, red) induced leukemia infiltration in BM and PB in U-937 AML xenografts (*N* = 7). (B) Immmunohistochemical analysis of the epiphyseal bone of U-937 AML xenografts stained with anti-hCD45 antibody indicate that CD81 overexpression (CD81+, left panel) induced higher BM (green outline) and adjacent muscle tissue AML infiltration (red outline) compared to control (CD81-, right panel). (C) Kaplan-Meier plot illustrating worse survival of mice xenografted with U937 CD81 overexpressing cells (CD81+) compared to control (*N* = 12). (D) In contrast, CD81 silencing in OCI-AML3 cells (CD81-, blue) reduced leukemia infiltration in BM, PB and spleen of AML xenografts (*N* = 7) and (E) prolonged survival in HNT-34 compared to control (*N* = 10). \**P* <0.05, \*\**P* <0.005, \*\*\**P* <0.0005.

Altogether, these results indicate that CD81 is an important determinant of leukemia chemoresistance and aggressiveness.

### CD81 is associated with LSC function in primary AML

As LSC are considered the main source of relapse^45–47^ and given the increased CD81 expression in blasts at the time of relapse, we investigated whether CD81 also affects LSC function. First, we assessed its expression within the AML LSC subpopulation, defined as CD34+/CD38-/CD90-/CD123+. This showed that CD81 was expressed within this LSC population and that CD81+ LSC fraction had increased at relapse when compared to samples obtained at diagnosis (Figure 5A) or between paired relapse and diagnosis samples (49% *vs.* 20%; Figure 5B). Similar results were found for the non-HSPC fraction defined as CD34+/CD38-/CD90-(53% *vs.* 8%; Supplemental Figure 5A, B). Then we investigated whether a high proportion of LSC positive for CD81 negatively affects AML outcome. Our results showed that non-CBF AML patients with a higher proportion of CD81+LSC at diagnosis were at increased risk of relapse and death (RFS: HR [CI] = 2.01 [1.16-3.48]; OS: HR [CI] = 2.38 [1.32-4.29]; Figure 5C). Interestingly, the fraction of LSC positive for CD81 was correlated with bulk CD81 protein expression (Figure 5D), suggesting that bulk CD81 expression likely reflects the proportion of LSC positive for CD81.

**Figure 5.**
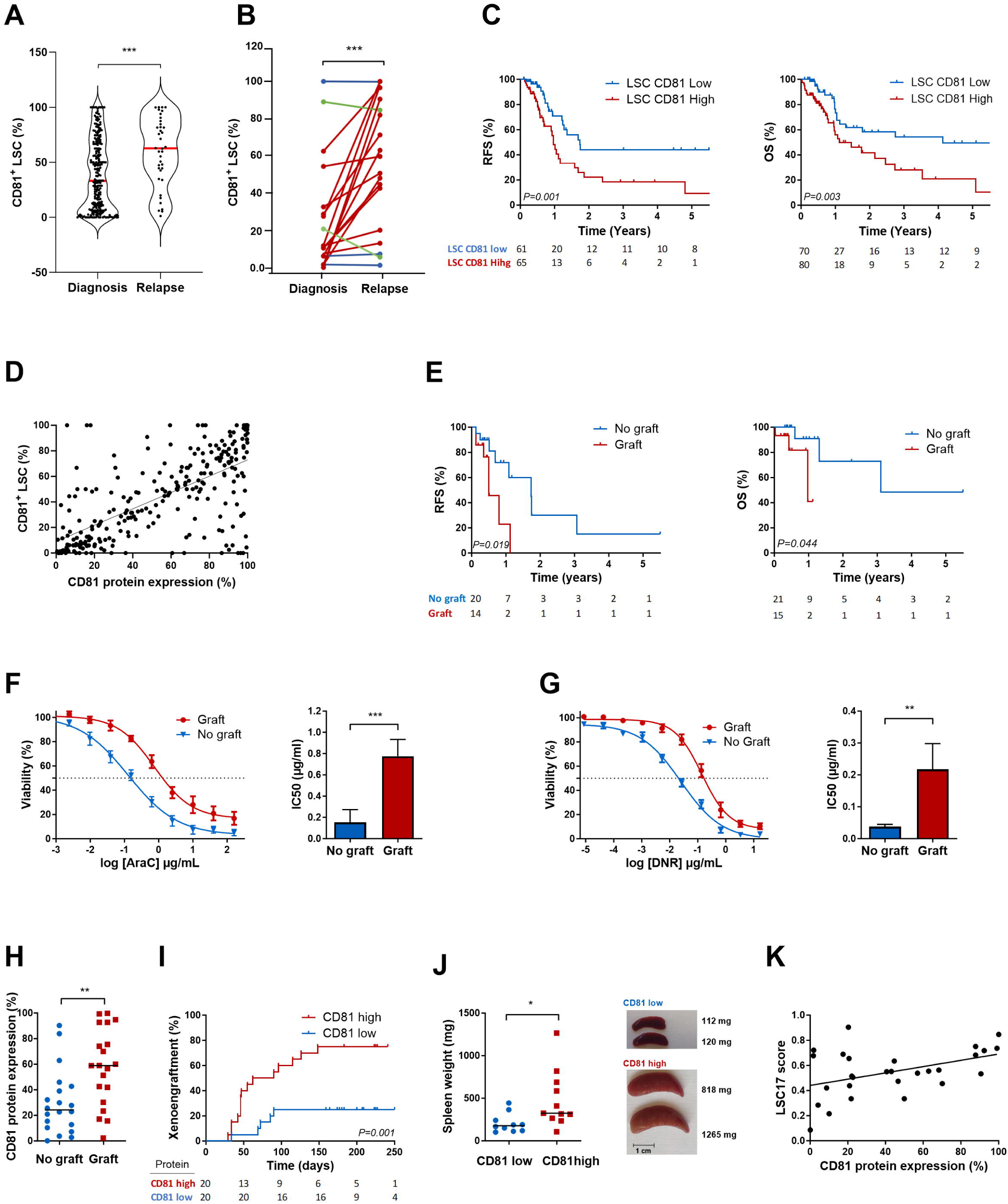
CD81 overexpression in AML is associated to increased LSC function, chemoresistance and worse clinical prognosis. (A) Violin plot indicating that the LSC fraction with CD81 protein expression increased from diagnosis to relapse (LSC defined as CD34+CD38-CD90-CD123+; *N* = 290) and (B) similar was found for paired diagnosis and relapse samples (red: increase, blue: no change, green: decrease; *N* = 19, paired t-test). (C) Kaplan-Meier plot showing worse RFS and OS in AML patients with higher CD81 (red) compared to lower CD81 (blue) within the LSC fraction at AML diagnosis (*N* = 150). (D) Scatterplot correlating CD81 protein expression within the LSC compartment with CD81 expression of the leukemia bulk (*N* = 290; *R*^2^= 0.44, *P* <0.0001). (E) Kaplan-Meier curves of RFS and OS for patients stratified by xenoengraftment capacity of primary AML (*N* = 56), where patients with engrafting AML (red) had worse outcome compared to those whose AML cannot engraft (blue). (F) Cytarabine-(*N* = 11) and (G) daunorubicin (*N* = 14) drug-response curves and bar plots of IC_50_ tested at AML diagnosis indicating higher resistance in engrafting (red) *vs*. not engrafting (blue) AML (MWU, Supplemental Figure 9). (H) High CD81 protein expression is not only associated with positive xenoengraftment (*N* = 40, MWU), but was also more successful as show by (I) cumulative incidence stratified by median expression (*N* = 40). (J) Scatter plot of spleen weights of xenografts comparing high *vs*. low median CD81 demonstrating that higher CD81 expression resulted in higher splenic infiltration (*N* = 21, MWU). (K) Scatterplot correlating LSC17 gene score and CD81 protein expression in non-CBF AML (*N* = 28. *R*^2^ = 0.19, *P* = 0.018). \**P* <0.05, \*\**P* <0.005, \*\*\**P* <0.0005.

To further characterize the relationship between CD81 overexpression and LSC function, we performed serial AML xenoengraftment experiments using immunocompromised NSG mice. As expected, patients whose AML xenoengrafted had worse outcome than those whose AML failed to engraft (RFS: HR [CI] = 4.45 [1.31-15.16]; OS: HR [CI] = 2.74 [1.15-6.51]; Figure 5E) and were more resistant to cytarabine and daunorubicin (Figure 5F-G). Moreover, our results also showed that AML with higher CD81 expression were not only associated with positive engraftment (Figure 5H) but also more successfully engrafted over time compared to low CD81 expressing AML (HR [CI] = 4.53 [1.64-12.56]; Figure 5I). Interestingly, PDX of AML with higher CD81 expression had larger spleens at sacrifice than those with low CD81 expression (Figure 5J), suggestive of higher extra-medullary infiltrative potential. Importantly, CD81 expression in non-CBF AML was correlated with the LSC17 gene signature score (Figure 2K), a known and reliable marker of functional LSC in AML.^34,48^

Taken together, these results unambiguously showed that high CD81 expression in primary AML is associated with functional LSC.

### CD81-targeting as a therapeutic strategy in AML

Having established the deleterious role of CD81 in AML and its translational relevance, we then investigated the therapeutic potential of an anti-hCD81 antibody. Firstly, we evaluated whether *in vitro* exposure of AML cells with this immunotherapy is able to inhibit their engraftment into NSG mice. Consistent with the data obtained using CD81-depleted cell lines, antibody-treatment of OCI-AML3 cells effectively reduced their engraftment into murine BM and PB and similarly reduced spleen infiltration, the preferential infiltration site of OCI-AML3 cells (Figure 6A). Additionally, targeting of CD81 using this therapeutic approach also increased *in vitro* sensitivity of OCI-AML3 to daunorubicin and cytarabine compared to IgG control (Figure 6B). Then, we evaluated potential toxicity by testing the *in vitro* effect of anti-hCD81 antibody on the viability of primary mononucleated BM cells (BMNC) obtained from a healthy individual. Of great interest, antibody-treatment neither affected BMNC viability at 4 and 24h nor altered BMNC cell cycle when compared to IgG control (Figure 6C-D). More importantly, no sign of BMNC toxicity was detected for the hematopoietic progenitors CFU-GEMM, CFU-GM, BFU-E or CFU-E (Figure 5E). Finally, we applied a curative protocol (Figure 6F) consisting of IP administration of either the anti-hCD81 antibody or control before, during and after intensive chemotherapy^29^ in leukemia bearing NSG mice using not only OCI-AML3 cells (Figure 6G), but also newly diagnosed primary AML cells (Figure 6H) and assessed treatment response by *in vivo* imaging and FCM, respectively. Our results showed that immunotargeting of CD81 strongly reduced tumor burden as early as one week after the last injection of the antileukemic drugs (Figure 6G, Supplemental Figure 6A) when using OCI-AML3 cells and more importantly, significantly reduced tumor burden after ten weeks off chemotherapy when using primary AML cells (Figure 6H, Supplemental Figure 6B). Moreover, in both cases, leukemia bearing mice treated with the anti-hCD81 antibody exhibited a less aggressive disease with delayed relapse and subsequently improved survival (HR [CI] = 2.60 [1.06-6.34], Figure 6I and HR PB-relapse [CI] = 5.228 [0.874-31.27], HR [CI] = 5.20 [0.53-50.6], Figure 6J).

**Figure 6.**
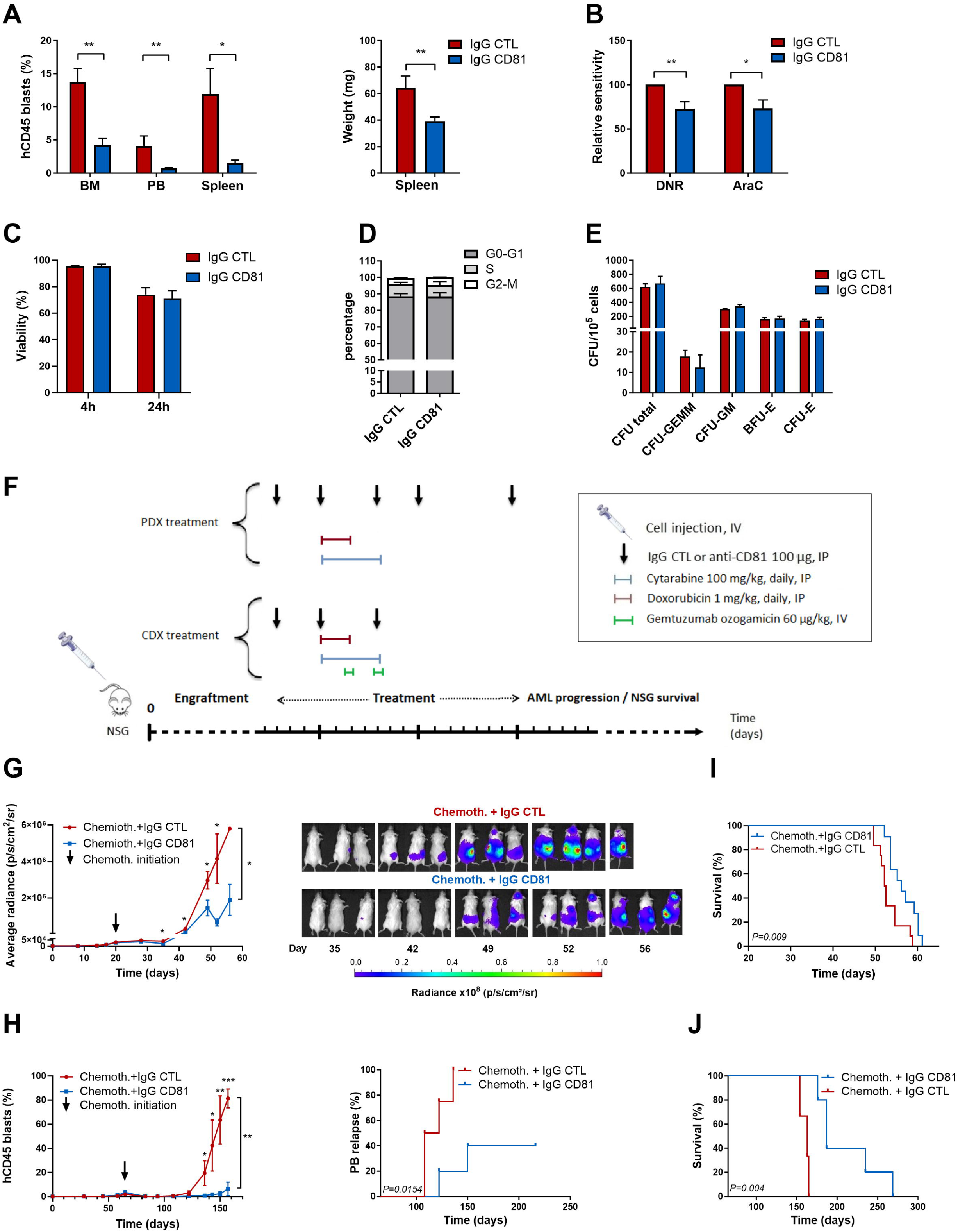
Efficacy and toxicity of therapeutic targeting of CD81 in AML. (A) Anti-hCD81 antibody-treatment decreased OCI-AML3 cell engraftment in BM, PB and spleen (N >7) and (B) sensitized OCI-AML3 cells to chemotherapies: daunorubicin (DNR, *N* = 6) and cytarabine (AraC, *N* = 4) compared to control. Anti-hCD81 antibody-treatment of normal BMNC cells affected neither viability (C), cell cycle (D) nor hematopoietic progenitor CFU (E) when compared to control. (F) Curative treatment protocols of OCI-AML3 CDX and PDX assessing anti-hCD81 antibody-treatment efficacy. When AML engraftment was 1-5% in PB, treatment was initiated with anti-hCD81 or control (black arrows) and chemotherapy consisting of cytarabine (5 days, blue), doxorubicin (3 days, dark red) and gemtuzumab ozogamicin for CDX (day 3 and 5, green). (G) Line graph (left panel) illustrates therapeutic efficacy of anti-CD81 antibody (blue, *N* = 11) in reducing tumor burden compared to control (red, *N* = 10) using OCI-AML3 CDX and corresponding *in vivo* images (right panel). (H) Line graph illustrating reduced AML burden in PB (left panel) and lower cumulative incidence of relapse (right panel) of PDX mice treated with anti-hCD81 antibody (blue, *N* = 5) compared to control (red, *N* = 3). Kaplan-Meier plots of (I) CDX- and (J) PDX mice stratified by treatment showing that prolonged OS with anti-hCD81 antibody *vs*. control. CTL: control, CFU: colony-forming unit, \**P* <0.01, \*\**P* <0.001, \*\*\**P* <0.0001.

In summary, immunotargeting of CD81 may represent a novel safe and effective therapeutic strategy for AML.

## Discussion

AML is a hematological stem cell malignancy originating from a rare subpopulation of LSC that is responsible for the accumulation of undifferentiated myeloid blast cells at the expense of the hematopoietic system.^49^ Recent sequencing studies have revealed that AML is genetically complex with patients frequently harboring many distinct driver mutations whereas no commonly dysregulated oncogenic pathways has been identified yet.^50^ This implies that specific targeting of any particular leukemogenic molecular alteration may not only be effective in just a small subset of patients but may also only eliminate a fraction of neoplastic cells, as multiple mutations frequently coexist within a leukemia. Therefore, it is increasingly evident that achieving long-term remissions or even curing AML would unlikely rely on a single targeted-therapy^51–53^ but instead should ideally require the development of treatment strategies effective against LSC and across a wide range of AML subtypes, irrespective of their mutational status. In this regard, AML-specific cell surface proteins detectable in most patients have raised particular theranostic interest such as CD33, CD123, CLL-1, CD44, CD47 or TIM-3 and are currently under investigation.^11,49^ Nevertheless, targeting these molecules is lagging behind expectations as severe toxicities caused by the lack of expression specificity and resistance frequently occur. For example, targeting CD33 (Siglec-3), a myeloid cell surface antigen highly expressed in most AML subtypes and LSC-enriched populations, using a recombinant humanized anti-hCD33 antibody conjugated to the cytotoxic calicheamicin (gemtuzumab ozogamicin)^54^ has shown efficacy especially in favorable AML while inducing severe toxicity given its expression on normal hematopoietic cells.

In this study, we aimed to further assess the therapeutic potential of CD81 in AML, a cell surface glycoprotein expressed on immune cells and known to be essential for B cell maturation and differentiation.^55,56^ Our results notably showed that CD81 is an independent prognostic marker of treatment response in AML, especially in non-favorable AML. Consistent with this, we functionally validated its deleterious role using gain and loss of function *in vitro* and *in vivo* approaches by demonstrating that CD81 expression affects leukemia aggressiveness and tumor burden by enhancing cell adhesion, migration and invasion processes.^57–59^ Indeed, tetraspanins such as CD81 are major components of specialized membrane microdomains known to interact with the extracellular matrix to mediate cell adhesion and motility,^24,60^ but a functional role of CD81 in myeloid cell adhesion has not been reported yet.

As LSC are considered to play a pivotal role in AML relapse, eradicating these cells by identifying a highly expressed if not specific therapeutic target remains critical to ultimately cure AML while sparing other important cell populations such as normal HSPC clones.^61^ As cell adhesion and motility are fundamental for LSC interaction with the BM niche microenvironment and given our results showing the importance of CD81 in these processes, we reasoned that this cell surface protein may also influence LSC function. For this, we first showed that CD81 is expressed within the AML-LSC-containing subpopulation commonly defined as CD34+CD38-CD90-. Indeed, previous studies combining immunophenotyping of AML samples and xenotransplantation have established that LSC are characterized by the CD34+CD38-CD90-immunophenotype in more than 90% of AML cases.^19,47^ Interestingly, CD81 was also expressed in the CD34+CD38-CD90-CD123+ subset which included CD123 (IL-3Rα), a well-established LSC specific cell surface marker.^12,13,62^ As targeting several antigens may be necessary to eliminate leukemic cells that persist at remission, combinatorial immunotargeting^63^ of both CD81 and CD123 may represent a novel therapeutic approach especially to circumvent antigen escape, a major cause of resistance to immunotherapy. Nevertheless, whether CD81 expression persists in LSC and progenitor populations of AML patients during clinical remission remains to be investigated. Consistent with previous studies related to CD33^64^ and CD123,^65^ we showed that CD81 was overexpressed on bulk tumor and LSC in most AML patients across different subtypes of AML, and more importantly, that its expression increased from diagnosis to relapse, suggesting a role for CD81 in AML treatment escape and disease progression. Therefore, targeting CD81 may not only be effective against persistent leukemic cells but also as second-line treatment when relapse occurs. Lastly, we demonstrated that CD81 expression is associated with functional LSC^9^ using serial xenoengraftment experiments as well as the LSC17 score,^48^ a known surrogate marker of LSC activity.

Finally, as targeting cell surface proteins are often associated with severe toxicity related to their expression in hematopoietic tissues,^12,47^ we evaluated whether anti-hCD81 antibody-treatment is a safe strategy. When exposed to normal BMNC, antibody-treatment revealed no hematopoietic toxicity neither alone nor in combination with intensive chemotherapy and was well tolerated by xenoengrafted mice. This is consistent with previous reports showing that (i) CD81 expression is low in normal BM cells,^32^ (ii) anti-mCD81 antibody-treatment is well-tolerated in inflammatory bowel diseased mice^66^ and more importantly, (iii) an anti-hCD81 antibody developed as an inhibitor of HCV infection^67^ and recognizing both human and monkey CD81 is well-tolerated in primates using several therapeutic doses over a prolonged time.^68^ Lastly, murine knockout models of CD81 have been established and no life threatening hematopoietic defects were observed except reduced B cell activity,^55,69,70^ mirroring the phenotype characterizing human autosomal recessive CD81 deficiency.^71^

In conclusion, we provide proof of concept for CD81 as a new relevant therapeutic target and diagnostic in AML. In particular, immunotargeting of CD81 combined with standard chemotherapy may represent a novel efficacious treatment strategy to improve AML outcome especially during post-remission therapy and at relapse given the importance of CD81 for LSC function.

## Acknowledgements

French National Cancer Institute INCa grants: SIRIC Oncolille 2014-17 (CP, MC), PRTK 2016 (CP, MC) and PLBio 2018 (MC), the Foundations Laurette Fugain 2012 (MC) and ARC 2014 (MC) and the Association Ligue contre le cancer 2016 (MC). The authors are grateful for the participation of the patients, and clinical and administrative staff implicated in this study.

## Authors’ contributions

FG, PP, NP, ND, MC wrote the manuscript; FG, PP, TB, SG, AB, FS, FD, AP, CR, MC performed experiments and analyzed results, CP, MC designed and financed experiments, CR, MC interpreted and discussed results, and CB is the referring physician of these patients.

## References

1. Visser, O. et al. Incidence, survival and prevalence of myeloid malignancies in Europe. European Journal of Cancer 48, 3257–3266 (2012).

2. Tiong, I. S. & Wei, A. H. New drugs creating new challenges in acute myeloid leukemia. *Genes*, Chromosomes and Cancer 58, 903–914 (2019).

3. Döhner, H. et al. Diagnosis and management of AML in adults: 2017 ELN recommendations from an international expert panel. Blood 129, 424–447 (2017).

4. Röllig, C. et al. Long-term prognosis of acute myeloid leukemia according to the new genetic risk classification of the European LeukemiaNet recommendations: evaluation of the proposed reporting system. J Clin Oncol 29, 2758–2765 (2011).

5. Dombret, H. & Gardin, C. An update of current treatments for adult acute myeloid leukemia. Blood 127, 53–61 (2016).

6. Vardiman, J. W. et al. The 2008 revision of the World Health Organization (WHO) classification of myeloid neoplasms and acute leukemia: rationale and important changes. Blood 114, 937–951 (2009).

7. Thol, F., Schlenk, R. F., Heuser, M. & Ganser, A. How I treat refractory and early relapsed acute myeloid leukemia. Blood 126, 319–327 (2015).

8. O’Reilly, E., Zeinabad, H. A. & Szegezdi, E. Hematopoietic versus leukemic stem cell quiescence: Challenges and therapeutic opportunities. Blood Reviews 50, 100850 (2021).

9. Lapidot, T. et al. A cell initiating human acute myeloid leukaemia after transplantation into SCID mice. Nature 367, 645–648 (1994).

10. Shlush, L. I. et al. Tracing the origins of relapse in acute myeloid leukaemia to stem cells. Nature (2017) doi:10.1038/nature22993.

11. Pollyea, D. A. & Jordan, C. T. Therapeutic targeting of acute myeloid leukemia stem cells. Blood 129, 1627–1635 (2017).

12. Taussig, D. C. Hematopoietic stem cells express multiple myeloid markers: implications for the origin and targeted therapy of acute myeloid leukemia. Blood 106, 4086–4092 (2005).

13. Jordan, C. et al. The interleukin-3 receptor alpha chain is a unique marker for human acute myelogenous leukemia stem cells. Leukemia 14, 1777–1784 (2000).

14. Majeti, R. et al. CD47 Is an Adverse Prognostic Factor and Therapeutic Antibody Target on Human Acute Myeloid Leukemia Stem Cells. Cell 138, 286–299 (2009).

15. Jaiswal, S. et al. CD47 Is Upregulated on Circulating Hematopoietic Stem Cells and Leukemia Cells to Avoid Phagocytosis. Cell 138, 271–285 (2009).

16. Kikushige, Y. et al. TIM-3 Is a Promising Target to Selectively Kill Acute Myeloid Leukemia Stem Cells. Cell Stem Cell 7, 708–717 (2010).

17. van Rhenen, A. et al. The novel AML stem cell–associated antigen CLL-1 aids in discrimination between normal and leukemic stem cells. Blood 110, 2659–2666 (2007).

18. Moshaver, B. et al. Identification of a Small Subpopulation of Candidate Leukemia-Initiating Cells in the Side Population of Patients with Acute Myeloid Leukemia. Stem Cells 26, 3059–3067 (2008).

19. Goardon, N. et al. Coexistence of LMPP-like and GMP-like leukemia stem cells in acute myeloid leukemia. Cancer Cell 19, 138–152 (2011).

20. Blair, A., Hogge, D. E., Ailles, L. E., Lansdorp, P. M. & Sutherland, H. J. Lack of Expression of Thy-1 (CD90) on Acute Myeloid Leukemia Cells With Long-Term Proliferative Ability In Vitro and In Vivo. Blood 89, 3104–3112 (1997).

21. Oren, R., Takahashi, S., Doss, C., Levy, R. & Levy, S. TAPA-1, the target of an antiproliferative antibody, defines a new family of transmembrane proteins. Mol Cell Biol 10, 4007–15 (1990).

22. Shoham, T., Rajapaksa, R., Kuo, C.-C., Haimovich, J. & Levy, S. Building of the Tetraspanin Web: Distinct Structural Domains of CD81 Function in Different Cellular Compartments. Mol Cell Biol 26, 1373–1385 (2006).

23. Yáñez-Mó, M., Barreiro, O., Gordon-Alonso, M., Sala-Valdés, M. & Sánchez-Madrid, F. Tetraspanin-enriched microdomains: a functional unit in cell plasma membranes. Trends in Cell Biology 19, 434–446 (2009).

24. Hemler, M. E. Tetraspanin functions and associated microdomains. Nat Rev Mol Cell Biol 6, 801–811 (2005).

25. Umeda, R. et al. Structural insights into tetraspanin CD9 function. Nat Commun 11, 1606 (2020).

26. Zimmerman, B. et al. Crystal Structure of a Full-Length Human Tetraspanin Reveals a Cholesterol-Binding Pocket. Cell 167, 1041–1051.e11 (2016).

27. Susa, K. J., Seegar, T. C., Blacklow, S. C. & Kruse, A. C. A dynamic interaction between CD19 and the tetraspanin CD81 controls B cell co-receptor trafficking. eLife 9, e52337 (2020).

28. Bradbury, L. E., Kansas, G. S., Levy, S., Evans, R. L. & Tedder, T. F. The CD19/CD21 signal transducing complex of human B lymphocytes includes the target of antiproliferative antibody-1 and Leu-13 molecules. J. Immunol. 149, 2841–2850 (1992).

29. Vences-Catalán, F. et al. CD81 is a novel immunotherapeutic target for B cell lymphoma. J. Exp. Med. 216, 1497–1508 (2019).

30. Quagliano, A., Gopalakrishnapillai, A., Kolb, E. A. & Barwe, S. P. CD81 knockout promotes chemosensitivity and disrupts in vivo homing and engraftment in acute lymphoblastic leukemia. Blood Advances 4, 4393–4405 (2020).

31. Vences-Catalán, F. et al. Targeting the tetraspanin CD81 reduces cancer invasion and metastasis. Proc Natl Acad Sci U S A 118, (2021).

32. Boyer, T. et al. Tetraspanin CD81 is an adverse prognostic marker in acute myeloid leukemia. Oncotarget (2016) doi:10.18632/oncotarget.11481.

33. Boyer, T. et al. Flow Cytometry to Estimate Leukemia Stem Cells in Primary Acute Myeloid Leukemia and in Patient-derived-xenografts, at Diagnosis and Follow Up. J Vis Exp (2018) doi:10.3791/56976.

34. Duployez, N. et al. The stem cell-associated gene expression signature allows risk stratification in pediatric acute myeloid leukemia. Leukemia 33, 348–357 (2019).

35. Nibourel, O. et al. Copy-number analysis identified new prognostic marker in acute myeloid leukemia. Leukemia 31, 555–564 (2017).

36. Schneider, C. A., Rasband, W. S. & Eliceiri, K. W. NIH Image to ImageJ: 25 years of image analysis. Nat Methods 9, 671–675 (2012).

37. Zhang, C. C. et al. Gemtuzumab Ozogamicin (GO) Inclusion to Induction Chemotherapy Eliminates Leukemic Initiating Cells and Significantly Improves Survival in Mouse Models of Acute Myeloid Leukemia. Neoplasia 20, 1–11 (2018).

38. R Core Team. R: A language and environment for statistical computing. (2022).

39. Tyner, J. W. et al. Functional genomic landscape of acute myeloid leukaemia. Nature 562, 526–531 (2018).

40. Cerami, E. et al. The cBio cancer genomics portal: an open platform for exploring multidimensional cancer genomics data. Cancer Discov 2, 401–404 (2012).

41. Gao, J. et al. Integrative analysis of complex cancer genomics and clinical profiles using the cBioPortal. Sci Signal 6, pl1 (2013).

42. Saland, E. et al. A robust and rapid xenograft model to assess efficacy of chemotherapeutic agents for human acute myeloid leukemia. Blood Cancer J 5, e297 (2015).

43. Shultz, L. D. et al. Human lymphoid and myeloid cell development in NOD/LtSz-scid IL2R gamma null mice engrafted with mobilized human hemopoietic stem cells. J. Immunol 174, 6477–6489 (2005).

44. Dick, J. E. & Lapidot, T. Biology of Normal and Acute Myeloid Leukemia Stem Cells. International Journal of Hematology 82, 389–396 (2005).

45. Stelmach, P. & Trumpp, A. Leukemic stem cells and therapy resistance in acute myeloid leukemia. haematol 108, 353–366 (2023).

46. Vetrie, D., Helgason, G. V. & Copland, M. The leukaemia stem cell: similarities, differences and clinical prospects in CML and AML. Nat Rev Cancer 20, 158–173 (2020).

47. Thomas, D. & Majeti, R. Biology and relevance of human acute myeloid leukemia stem cells. Blood 129, 1577–1585 (2017).

48. Ng, S. W. K. et al. A 17-gene stemness score for rapid determination of risk in acute leukaemia. Nature 540, 433–437 (2016).

49. Mitchell, K. & Steidl, U. Targeting Immunophenotypic Markers on Leukemic Stem Cells: How Lessons from Current Approaches and Advances in the Leukemia Stem Cell (LSC) Model Can Inform Better Strategies for Treating Acute Myeloid Leukemia (AML). Cold Spring Harb Perspect Med 10, a036251 (2020).

50. Papaemmanuil, E. et al. Genomic Classification and Prognosis in Acute Myeloid Leukemia. N Engl J Med 374, 2209–2221 (2016).

51. Welch, J. S. et al. The Origin and Evolution of Mutations in Acute Myeloid Leukemia. Cell 150, 264–278 (2012).

52. Ding, L. et al. Clonal evolution in relapsed acute myeloid leukaemia revealed by whole-genome sequencing. Nature 481, 506–510 (2012).

53. Parkin, B. et al. Clonal evolution and devolution after chemotherapy in adult acute myelogenous leukemia. Blood 121, 369–377 (2013).

54. Bross, P. F. et al. Approval summary: gemtuzumab ozogamicin in relapsed acute myeloid leukemia. Clin Cancer Res 7, 1490–6 (2001).

55. Maecker, H. T. & Levy, S. Normal lymphocyte development but delayed humoral immune response in CD81-null mice. J Exp Med 185, 1505–10 (1997).

56. Vences-Catalán, F. et al. A mutation in the human tetraspanin CD81 gene is expressed as a truncated protein but does not enable CD19 maturation and cell surface expression. J. Clin. Immunol. 35, 254–263 (2015).

57. Paterson, E. K. & Courtneidge, S. A. Invadosomes are coming: new insights into function and disease relevance. FEBS J 285, 8–27 (2018).

58. Jacquemet, G., Hamidi, H. & Ivaska, J. Filopodia in cell adhesion, 3D migration and cancer cell invasion. Current Opinion in Cell Biology 36, 23–31 (2015).

59. Bari, R. et al. Tetraspanins regulate the protrusive activities of cell membrane. Biochemical and Biophysical Research Communications 415, 619–626 (2011).

60. Termini, C. M. & Gillette, J. M. Tetraspanins Function as Regulators of Cellular Signaling. Front. Cell Dev. Biol. 5, (2017).

61. Baum, C. M., Weissman, I. L., Tsukamoto, A. S., Buckle, A. M. & Peault, B. Isolation of a candidate human hematopoietic stem-cell population. Proc. Natl. Acad. Sci. U.S.A. 89, 2804–2808 (1992).

62. Vergez, F. et al. High levels of CD34+CD38low/-CD123+ blasts are predictive of an adverse outcome in acute myeloid leukemia: a Groupe Ouest-Est des Leucemies Aigues et Maladies du Sang (GOELAMS) study. Haematologica 96, 1792–8 (2011).

63. Haubner, S. et al. Coexpression profile of leukemic stem cell markers for combinatorial targeted therapy in AML. Leukemia 33, 64–74 (2019).

64. Baer, M. R. High frequency of immunophenotype changes in acute myeloid leukemia at relapse: implications for residual disease detection (Cancer and Leukemia Group B Study 8361). Blood 97, 3574–3580 (2001).

65. Bras, A. E. et al. CD123 expression levels in 846 acute leukemia patients based on standardized immunophenotyping. Cytometry 96, 134–142 (2019).

66. Hasezaki, T., Yoshima, T. & Mine, Y. Anti-CD81 antibodies reduce migration of activated T lymphocytes and attenuate mouse experimental colitis. Sci Rep 10, 6969 (2020).

67. Bailly, C. & Thuru, X. Targeting of Tetraspanin CD81 with Monoclonal Antibodies and Small Molecules to Combat Cancers and Viral Diseases. Cancers 15, 2186 (2023).

68. Vexler, V. et al. Target-mediated drug disposition and prolonged liver accumulation of a novel humanized anti-CD81 monoclonal antibody in cynomolgus monkeys. MAbs 5, 776– 786 (2013).

69. Miyazaki, T. Normal development but differentially altered proliferative responses of lymphocytes in mice lacking CD81. The EMBO Journal 16, 4217–4225 (1997).

70. Tsitsikov, E. N., Gutierrez-Ramos, J. C. & Geha, R. S. Impaired CD19 expression and signaling, enhanced antibody response to type II T independent antigen and reduction of B-1 cells in CD81-deficient mice. Proc. Natl. Acad. Sci. U.S.A. 94, 10844–10849 (1997).

71. Van Zelm, M. C. et al. CD81 gene defect in humans disrupts CD19 complex formation and leads to antibody deficiency. J. Clin. Invest. 120, 1265–1274 (2010).

