## Supplemental_Material for "Tetraspanin CD81 promotes leukemia stem cell function and represents a new therapeutic vulnerability in acute myeloid leukemia"

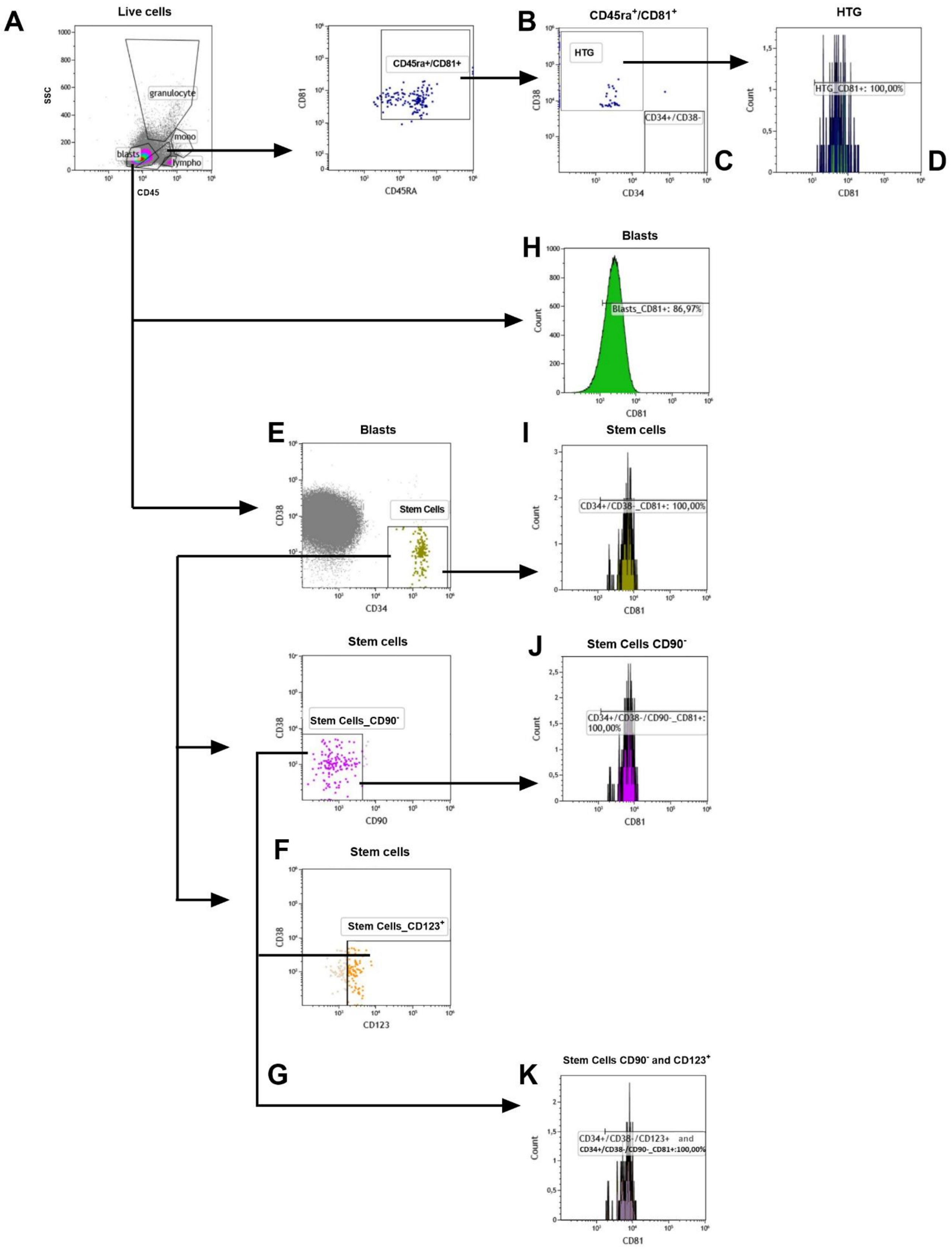

A
Live cells
SSC
granulocyte
mono
blasts
lympho
CD45
CD81
CD45ra+/CD81+
CD45RA
B
CD45ra+/CD81+
HTG
CD38
CD34+/CD38-
CD34
C
HTG
Count
HTG_CD81+: 100,00%
CD81
D
H
Blasts
Count
Blasts_CD81+: 86,97%
CD81
E
Blasts
Stem Cells
CD38
CD34
I
Stem cells
Count
CD34+/CD38-_CD81+: 100,00%
CD81
Stem cells
Stem Cells_CD90-
CD38
CD90
J
Stem Cells CD90-
Count
CD34+/CD38-/CD90-_CD81+: 100,00%
CD81
F
Stem cells
Stem Cells_CD123+
CD38
CD123
G
K
Stem Cells CD90- and CD123+
Count
CD34+/CD38-/CD123+ and CD34+/CD38-/CD90-_CD81+:100,00%
CD81

**Supplemental Figure 1** Flow cytometry (FCM) gating strategy of primary AML.

(A) Gating strategy using all bone marrow (BM) cells, where we first defined blasts (H) as $CD45^{dim}/SS^{low}$ and (B) hematogones as CD45RA+/CD81+/CD38+ [HTG].[1] (C-D) These gates were used to set the threshold for both CD38 and CD81, respectively.[2] (E-I) Stem cell population were gated as CD34+/CD38-[3] using the blast population. (F-J) Based on Stem cell population [CD34+/CD38-] engaged abnormal progenitor cells are defined as Stem Cells CD90- [CD34+/CD38-/ CD90-].[4,5] (F-G-K) Leukemia stem cells (LSC) were subsequently defined as Stem Cells CD90- and CD123+ [CD34+/CD38-/CD90-/CD123+].[6] CD81 expression was analyzed for the (H) blast-, (I) common stem cell-, (J) abnormal progenitor or non-HSPC and (K) leukemia stem cell (LSC) populations.

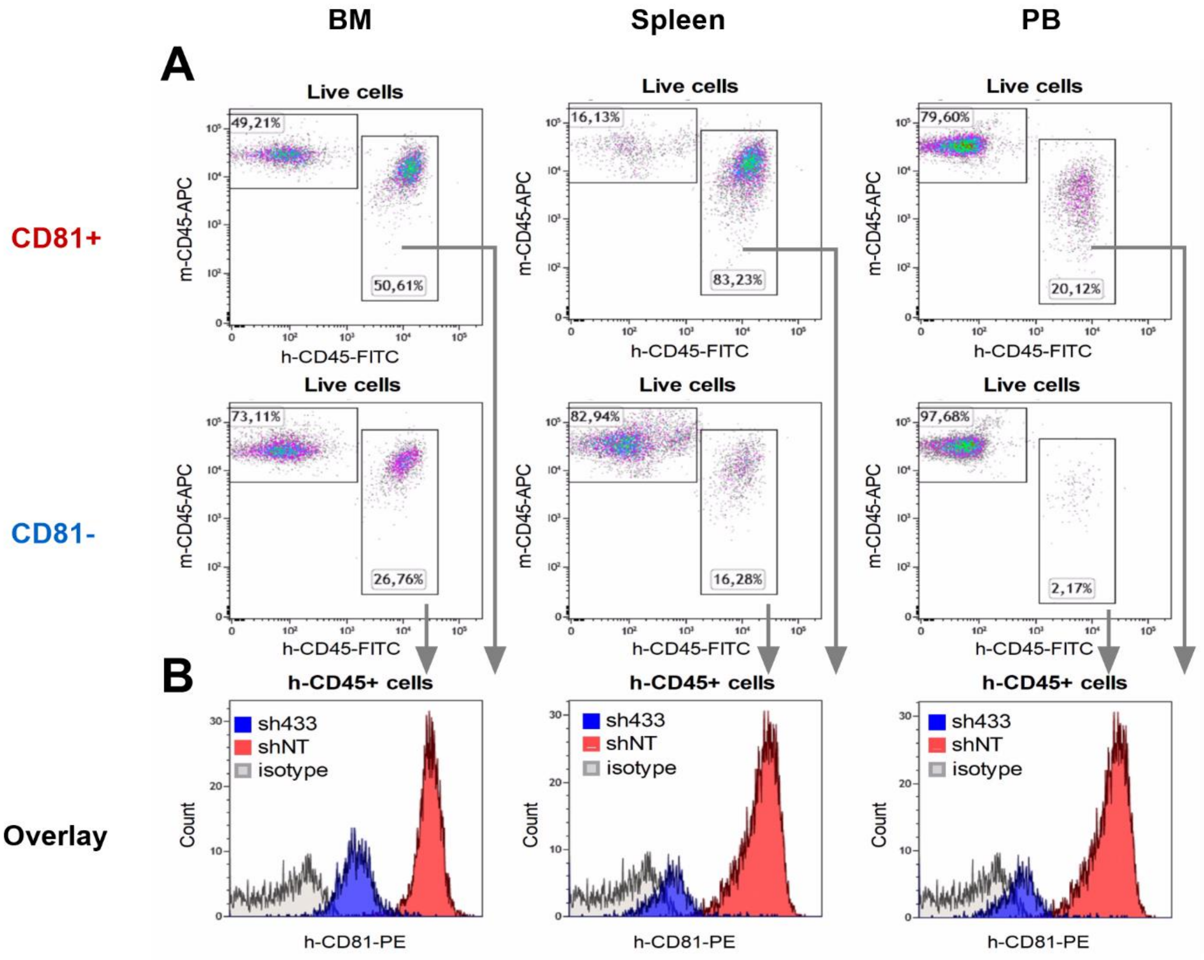


**Supplemental Figure 2** Engraftment and CD81 expression in *in vivo* models by flow cytometry (FCM).

Engraftment and CD81 expression was determined in bone marrow (BM), spleen and peripheral blood (PB) of CDX mice at sacrifice. (A) Gating strategy, we first defined AML blasts as [hCD45+/mCD45-] based on all live cells (FSC/SSC). (B) Overlay of CD81 expression using both CD81+ and CD81- model confirmed that no derivation of the phenotype had occurred during the *in vivo* experiment.

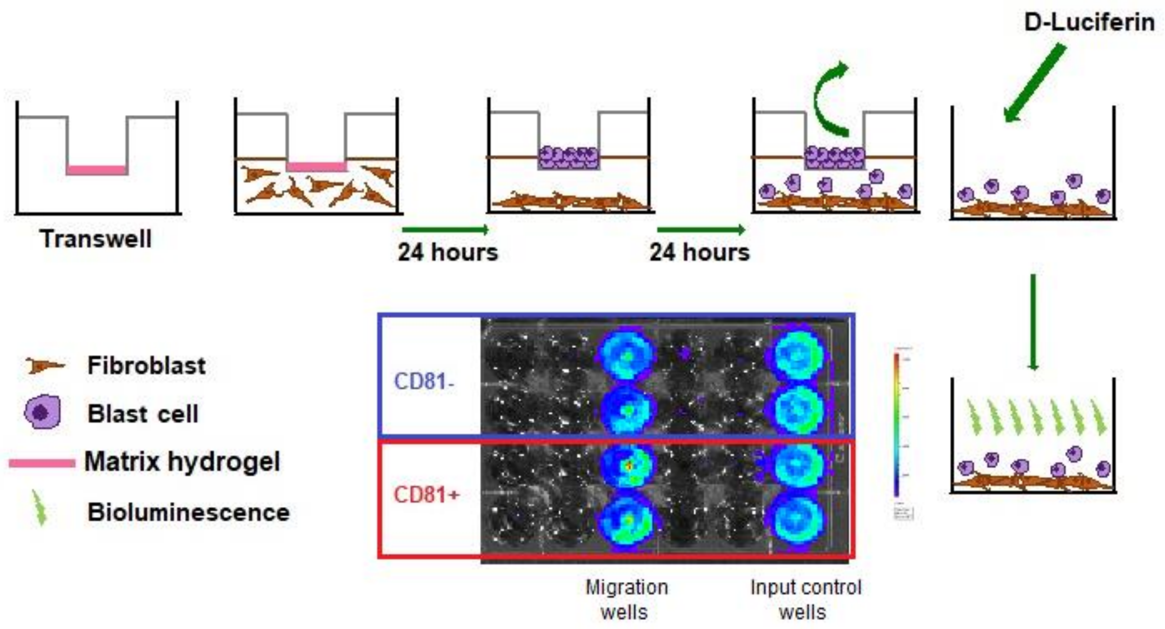


**Supplemental Figure 3** Schematic representation of Transwell migration and invasion assay. The migration assay was done without addition of hydrogel layer on top of the membrane, whereas the invasion assay included a matrix layer. In both cases the bottom well was coated with HS-5 bone marrow fibroblasts. Luminescence detection of cells passing to the bottom well vs. input control reflected respective migration and invasion capacity.

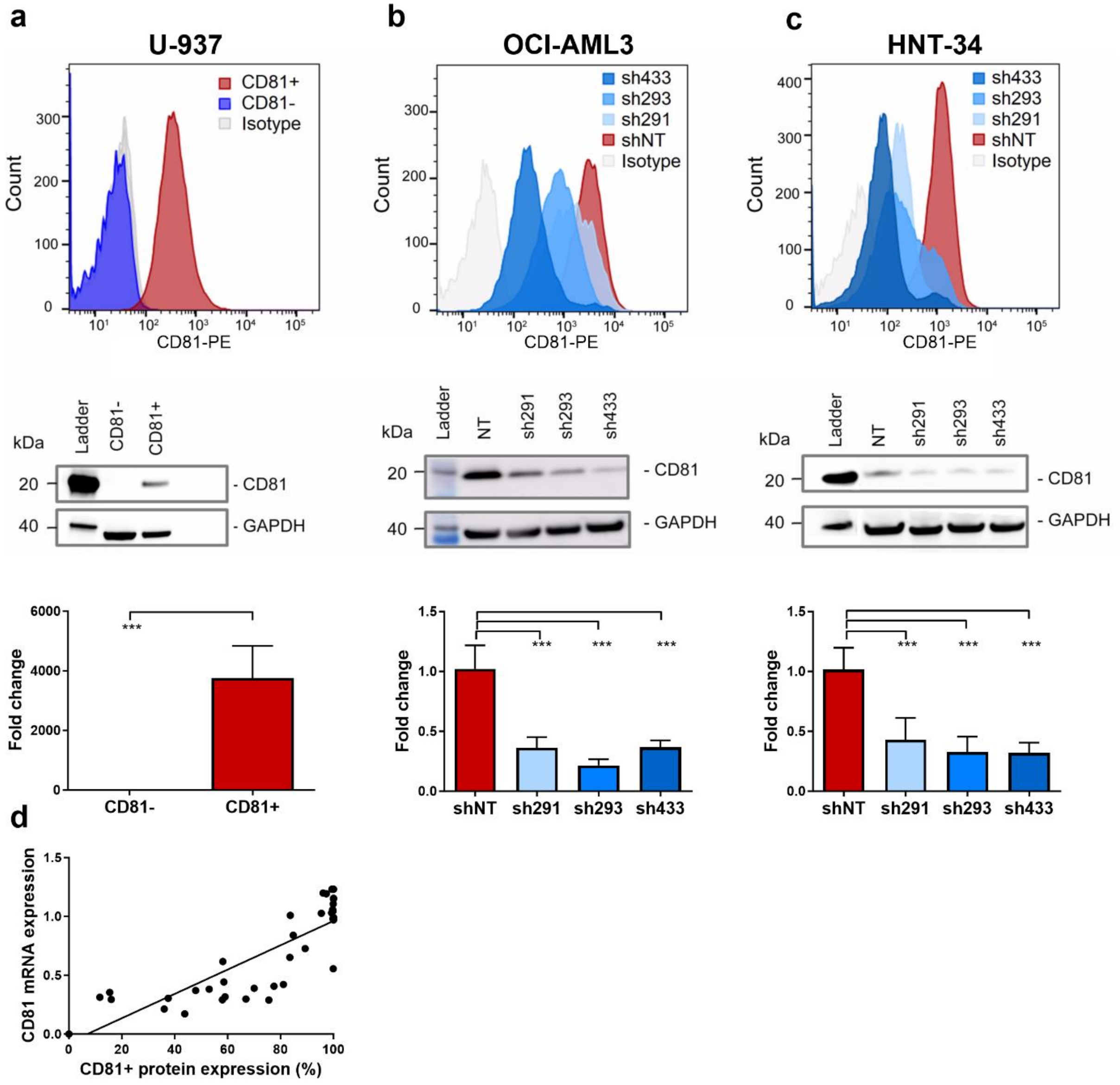


**Supplemental Figure 4** AML cell models with modified CD81 expression.

Shown are CD81 expression levels from both basal and modified variants of three different established AML cell lines (A) U-937, (B) OCI-AML3 and (C) HNT-34, quantified by FCM (top), Western blot (middle) and RT-PCR (bottom graphs). MWU-test, *** $P$ <0.0005. (D) Scatterplot showning correlation between CD81 protein and mRNA expression ($P$ = 0.010).

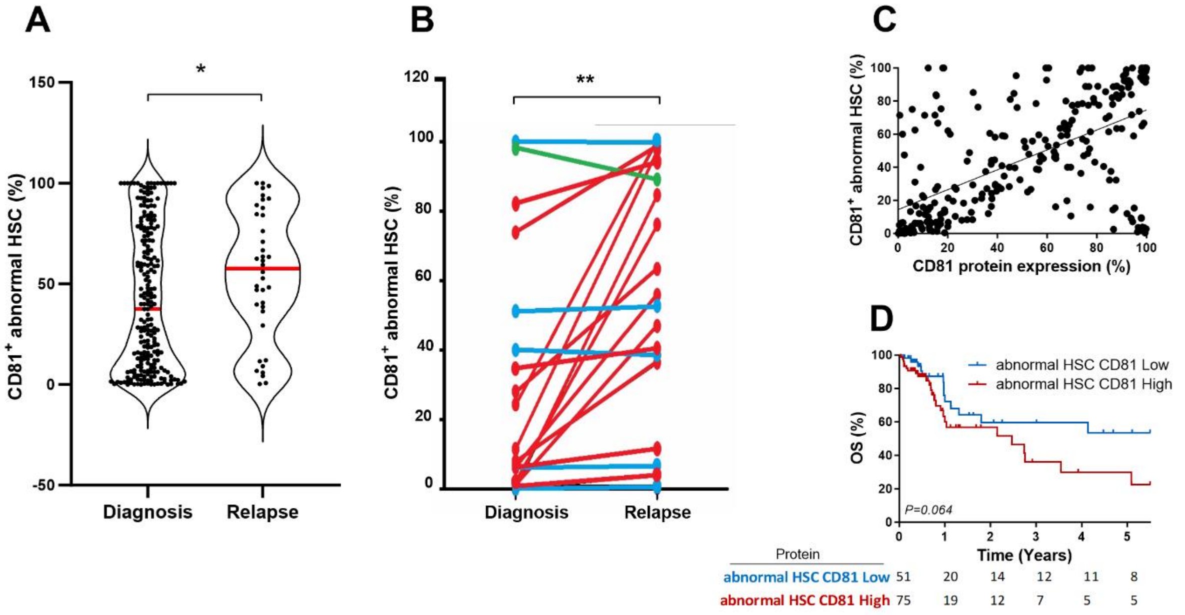


**Supplemental Figure 5** CD81 expression on the abnormal progenitor fraction.

(A) CD81 was expressed on abnormal HSPC defined as CD34+CD38-CD90- and this fraction increased from diagnosis to relapse and (B) in paired samples 53% vs. 8%, *P* = 0.0018 paired t test). Pairs are highlighted in red when the fraction increased, in green when decreased and in blue when unchanged from diagnosis to relapse. (C) CD81 protein expression on blasts was correlated to the CD81+ fraction of abnormal HSPC (*P* <0.0001). (D) Kaplan-Meier plot showing OS of patients with AML with higher vs lower median fraction of CD81+ abnormal HSPC CD34+CD38-CD90- (HR [95% CI] = 1.85 [0.98-3.5]).

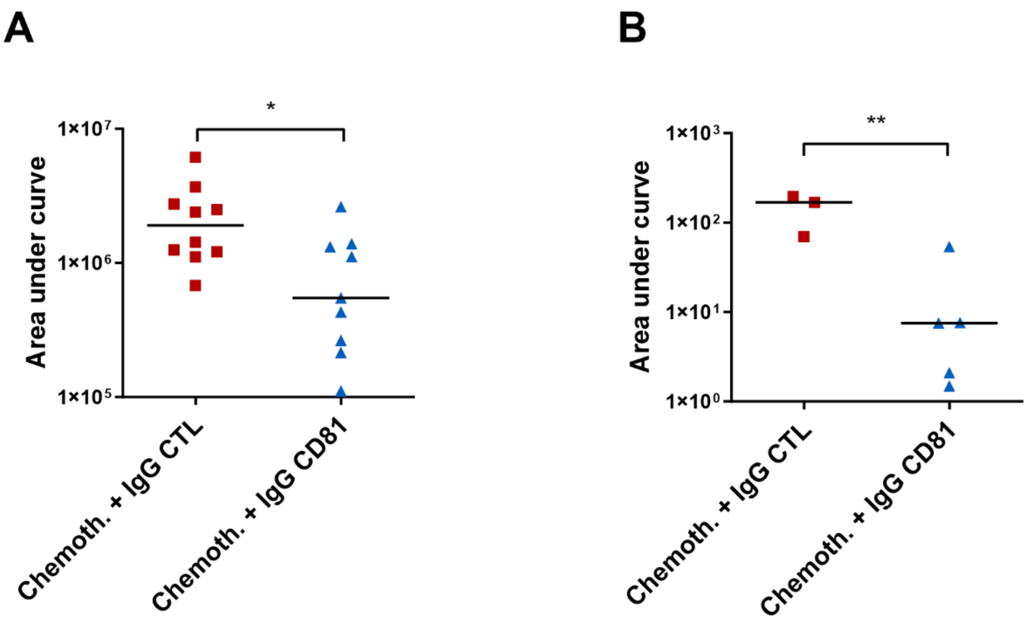


**Supplemental Figure 6** Therapeutic use of anti-hCD81 antibody on CDX and PDX.

(A) AUC comparison of OCI-AML3 CDX treated with anti-hCD81 antibody in association with chemotherapy (blue, *N* = 11) reduced leukemia burden measured by luminescence in CDX mice over a period of 8 weeks, compared to isotype control (red, *N* = 10, MWU).

(B) AUC of AML burden in PB of PDX mice quantified by FCM hCD45+ analysis from primary AML cell injection over a period of 22 weeks, comparing chemotherapy treatment combined with either anti-hCD81 antibody (blue, *N* = 5) or IgG control (red, *N* = 3).

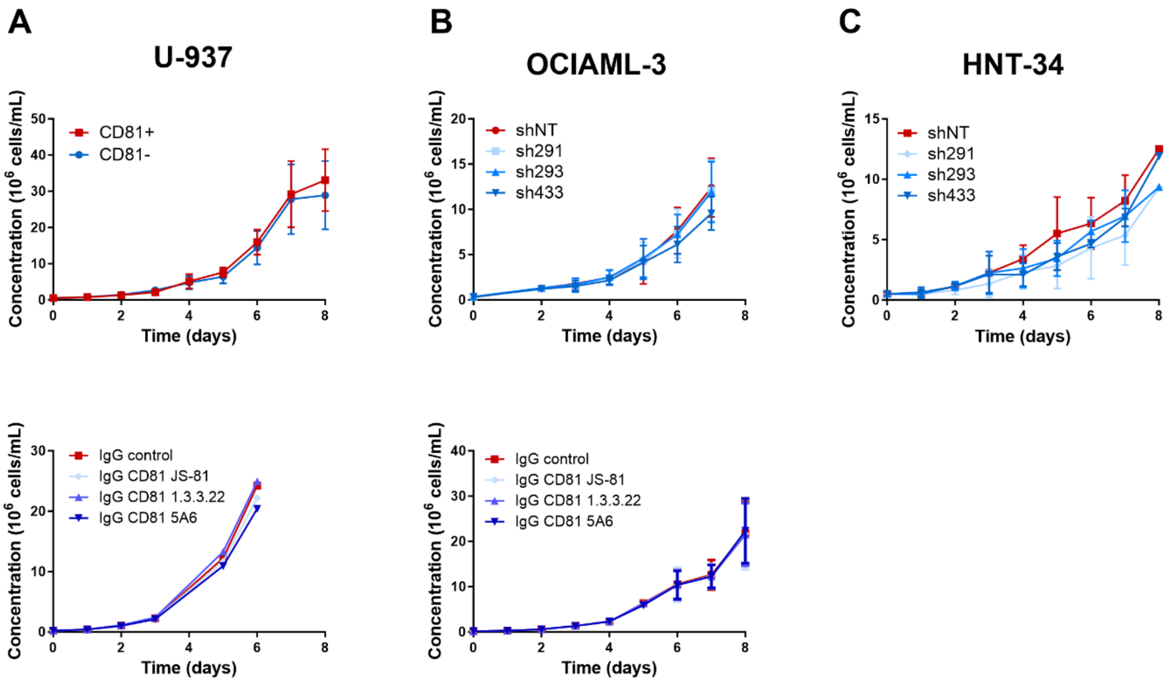


**Supplemental Figure 7** Proliferation of AML with modified CD81 expression and with anti-hCD81 antibody-treatment**.**

Neither AML model (A) U-937 (*N* = 6), (B) OCI-AML3 (*N* = 6) nor (C) HNT-34 cells (*N* = 5) showed altered proliferation following modulation of CD81 expression (top panel) or anti-hCD81 antibody treatment (bottom panel). Indicated cells (0.25 $\times 10^6$ cells/mL) were seeded into 6-well plates (Corning) and counted every 24 h by Cell Lab Quanta MPL Analyzer (Beckman Coulter) for 7 days. Treatment with three different anti-hCD81 antibodies has no effect on AML cell proliferation compared to isotype control.

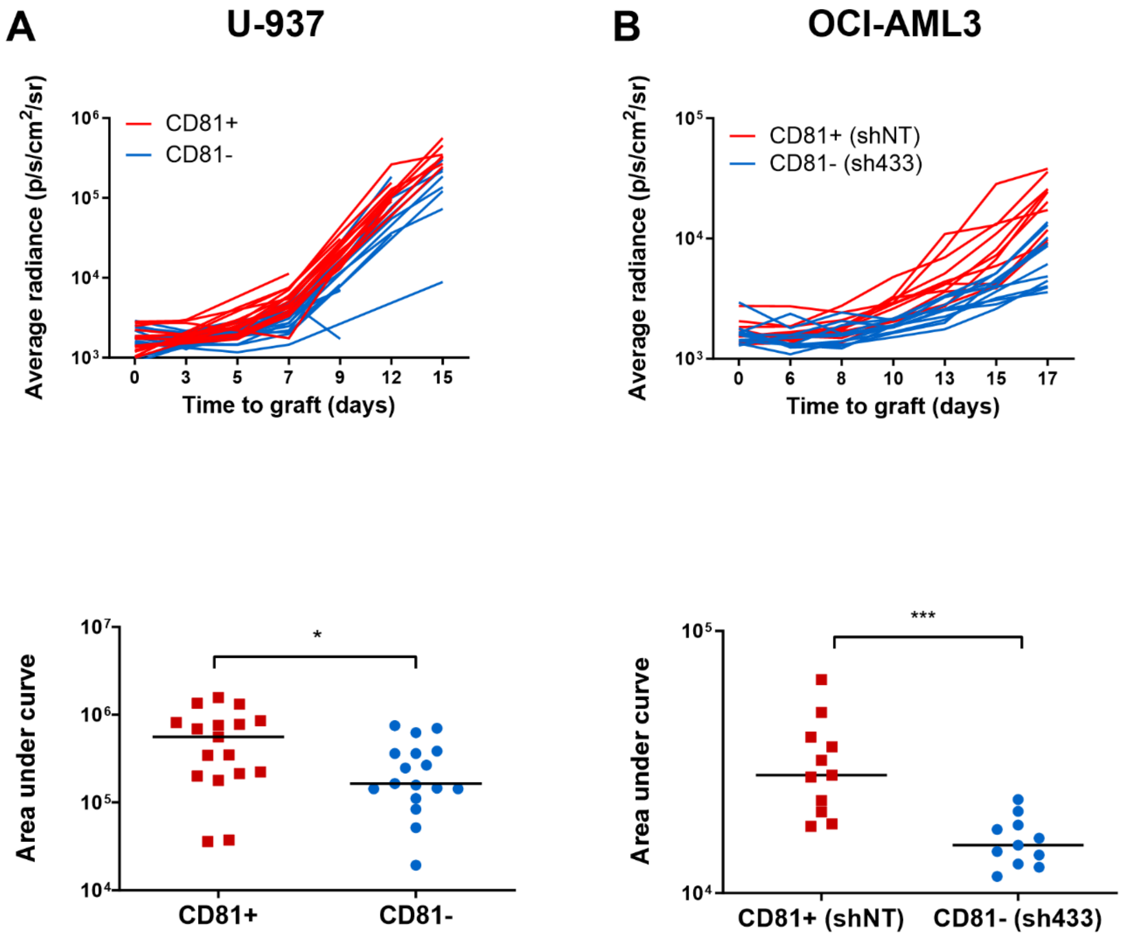


**Supplemental Figure 8** High CD81 expression increases xenoengraftment.

(A) Line graphs of each individual xenoengraftment, based on luminescence measured by *in vivo* imaging, shows increased engraftment rate of CD81+ (red) compared to CD81- (blue) U-937 cells, combined graphs are shown in main Figure 3A ($N$ = 17). Bottom: dot plot representing area under the curve (AUC) of each xenoengraftment grouped by CD81+ *vs.* CD81- (median: 5.61 x$10^5$ *vs.* 1.65 x$10^5$, $P$ = 0.030 MWU). (B) CD81 knockdown (blue) decreased xenoengraftment as shown for CD81 depleted OCI-AML3 cells compared to control (red). Individual xenografts are shown ($N$ = 11), corresponding to main Figure 3B with combined graphs. Bottom: dot plot representing AUC of each xenoengraftment grouped by control (red) *vs.* CD81 knockdown (blue, median = 2.82 x$10^4$ *vs*. 1.52 x$10^4$, $P$ = 0.0006 MWU). * $P$ <0.05; ** $P$ <0.005; *** $P$ <0.0005.

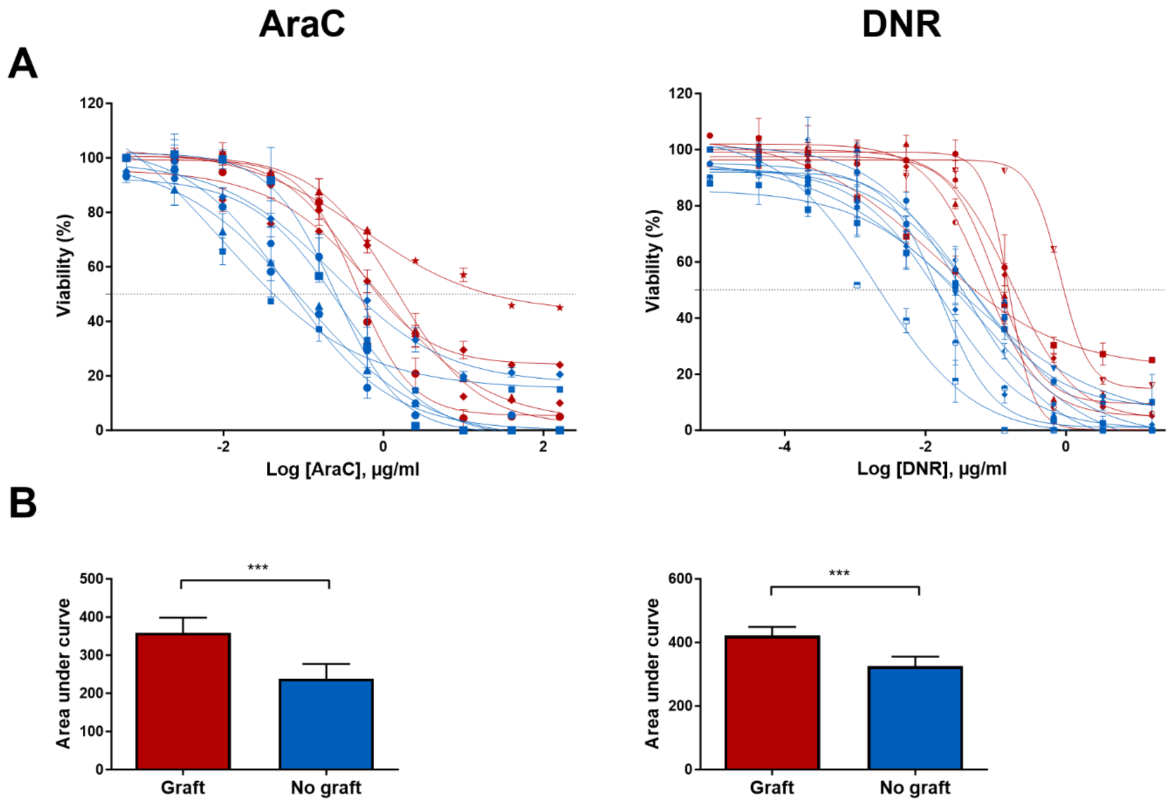


**Supplemental Figure 9** Chemoresistance of primary AML with xenoengraftment capacity.

(A) All individual drug-response curves for main Figures 2I and 2J for primary AML not engrafting (blue) and engrafting (red) demonstrating that latter AML were more resistant to cytarabine (AraC) and daunorubicin (DNR). Error bars indicate duplicate measurements. (B) Boxplots demonstrating the difference estimated as median AUC for AraC and DNR comparing primary AML not engrafting vs. engrafting. *** *P* <0.0005, MWU-test.

References


1. Muzzafar T, Medeiros LJ, Wang SA, et al. Aberrant underexpression of CD81 in precursor B-cell acute lymphoblastic leukemia: utility in detection of minimal residual disease by flow cytometry. *Am J Clin Pathol*. 2009;132(5):692–698.
2. Boyer T, Gonzales F, Plesa A, et al. Flow Cytometry to Estimate Leukemia Stem Cells in Primary Acute Myeloid Leukemia and in Patient-derived-xenografts, at Diagnosis and Follow Up. *JoVE*. 2018;(133):56976.
3. Bhatia M, Wang JC, Kapp U, Bonnet D, Dick JE. Purification of primitive human hematopoietic cells capable of repopulating immune-deficient mice. *Proc Natl Acad Sci U S A*. 1997;94(10):5320–5325.
4. Goardon N, Marchi E, Atzberger A, et al. Coexistence of LMPP-like and GMP-like leukemia stem cells in acute myeloid leukemia. *Cancer Cell*. 2011;19(1):138–152.
5. Majeti R, Park CY, Weissman IL. Identification of a hierarchy of multipotent hematopoietic progenitors in human cord blood. *Cell Stem Cell*. 2007;1(6):635–645.
6. Jordan CT, Upchurch D, Szilvassy SJ, et al. The interleukin-3 receptor alpha chain is a unique marker for human acute myelogenous leukemia stem cells. *Leukemia*. 2000;14(10):1777–84.
